# Chloride availability limits Slc4a9-dependent salivary secretion

**DOI:** 10.64898/2026.10.08.757782

**Authors:** Elías Vera-Sigüenza, Ramin Nashebi

**Affiliations:** Okinawa Institute of Science and Technology (OIST), Onna, Okinawa, Japan; School of Mathematics, University of Birmingham, Birmingham, England, United Kingdom

**Keywords:** salivary secretion, anion exchanger 4, intracellular chloride, chloride transport, acinar cell, mathematical modelling

## Abstract

Anion exchanger 4 (Ae4), encoded by solute carrier family 4 member 9 (Slc4a9), is required for normal salivary secretion, but how it supports fluid secretion remains poorly understood. Nearly a decade ago, we developed a mathematical model that attributed reduced secretion following Ae4 knockout primarily to Ae4 mediated monovalent cation transport.

Here, we reassess that interpretation by developing a dynamic model of salivary fluid secretion informed by subsequent experimental findings. The model incorporates *β*-adrenergic regulation of Ae4 and an additional apical chloride conductance shared by the control and knockout. To isolate the contribution of Ae4, the NKCC1 and Ae2 cycle rates calculated from the evolving control state are also prescribed in the knockout. Under these conditions, the knockout model secretes 20% less fluid over 600 s despite identical apical chloride conductances. The control begins with a larger intracellular chloride store and continues to acquire chloride through Ae4 during stimulation. The intracellular chloride difference increases during stimulation, whilst the weaker apical electrochemical driving force in the knockout reduces chloride export and fluid secretion. The analysis also identifies the alkalinity requirement of sustained Ae4 transport and quantifies the additional NKCC1 and Ae2 transport needed to compensate for its loss. These results support intracellular chloride availability as a mechanism linking Ae4 loss to reduced salivary secretion.

## 1 Introduction

Saliva production commences in the acinus, a cluster of acinar cells arranged around a central cavity known as the acinar lumen (Fig. 1). Ion transport across these cells establishes an osmotic gradient that draws water into the lumen, producing a fluid referred to as primary saliva (Melvin et al, 2005). As this initial secretion passes through the ductal system, its composition is modified prior to entering the oral cavity (Yusuke et al, 1973; Thaysen et al, 1954; Young and Schögel, 1966). The rate-limiting step in this process is the transcellular movement of chloride ions (Cl^−^). Acinar cells accumulate intracellular Cl^−^ at concentrations above those expected at electrochemical equilibrium, creating a pool available for secretion (Melvin et al, 2005; Peña-Münzenmayer et al, 2015, 2016; Catalán et al, 2025). Neurotransmitters such as acetylcholine (ACh) initiate a signalling cascade that ultimately raises intra-cellular calcium (Ca^2+^), which in turn leads to the activation of apical membrane Cl^−^ channels (Fig. 1). The subsequent efflux of Cl^−^ drives the parallel movement of sodium ions (Na^+^) and water, generating primary saliva (Melvin et al, 2005; Sneyd et al, 2021; Vera-Sigüenza et al, 2018).

**Fig. 1.**
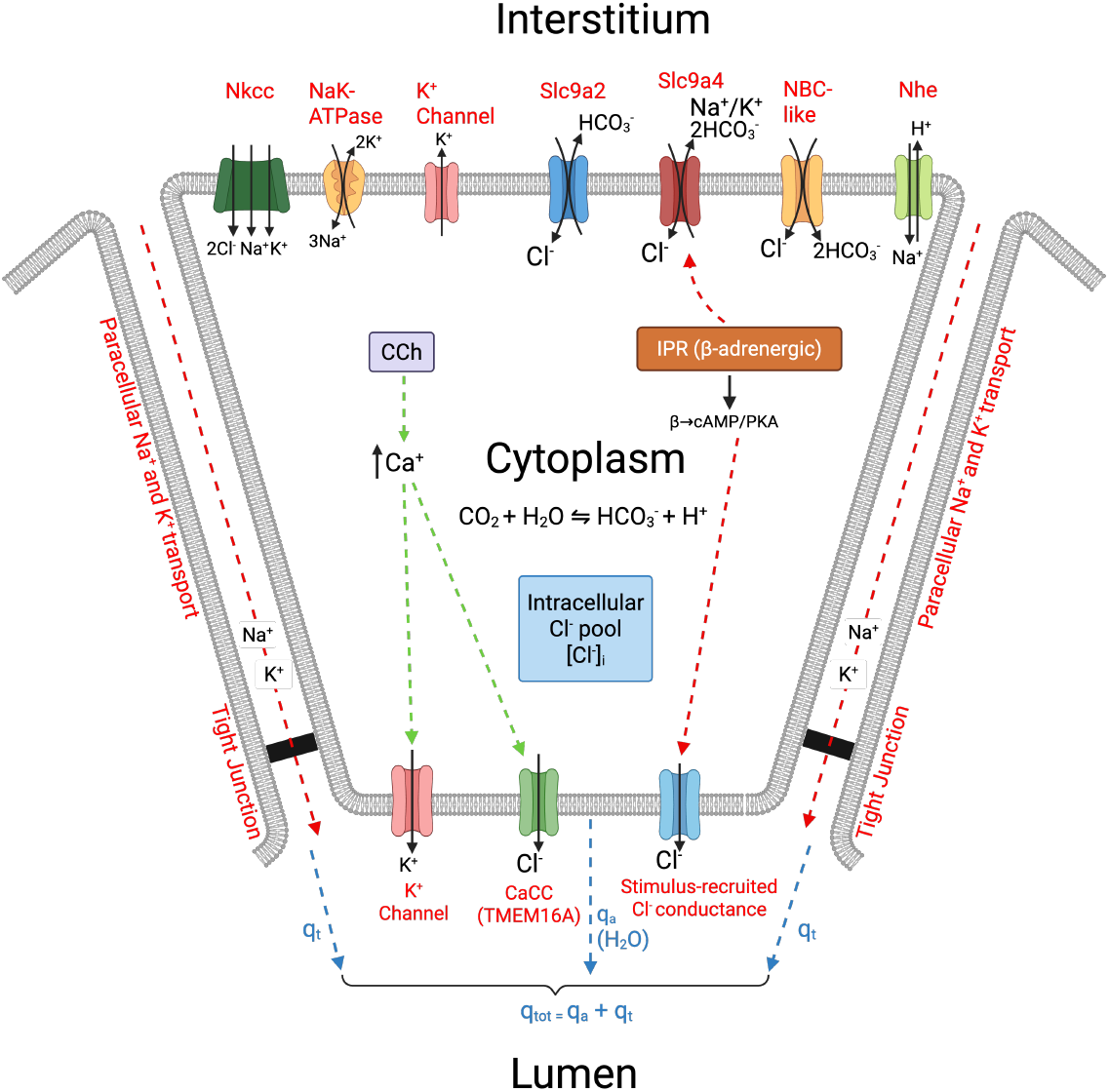
Schematic diagram of the salivary acinar cell model. Basolateral transport through NKCC1, Ae2, Ae4, NHE1, the Na^+^/K^+^ ATPase, and an NBCe1-B pathway recruited during stimulation regulates intracellular ion concentrations and replenishes the intracellular Cl^−^ pool. Cholinergic stimulation increases intracellular Ca^2+^ and activates apical Cl^−^ secretion through Tmem16A, whilst *β*-adrenergic stimulation acts through cAMP and PKA to activate Ae4 and an additional apical Cl^−^ conductance. Chloride secretion into the lumen generates the osmotic gradient that drives water flow and primary saliva formation. The model also includes paracellular 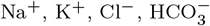, and water transport through the tight junctions.

Promoting water flow is not enough; the cell must also sustain it. To do so, acini must replenish their intracellular Cl^−^ by drawing from the interstitial space. The principal pathway for this uptake is the basolateral Na^+^-K^+^-Cl^−^ cotransporter (NKCC1), which imports 1 Na^+^, 1 potassium (K^+^), and 2 Cl^−^ ions into the cell. Pharmacological inhibition of NKCC1 sharply reduces acinar salivary secretion, whilst mice lacking NKCC1 exhibit an approximately 70% reduction in salivation (Martinez and Cassity, 1983; Evans et al, 2000). Yet neither pharmacological inhibition nor genetic deletion abolishes secretion completely (Peña-Münzenmayer et al, 2015). Additional Cl^−^ uptake pathways contribute to the secretory process (Nauntofte, 1992).

The molecular machinery underlying these additional pathways remained uncertain for decades (Melvin et al, 2005; Peña-Münzenmayer et al, 2015; Sneyd et al, 2021). Researchers proposed that one such route coupled Na^+^/H^+^ exchange through the Na^+^/H^+^ exchanger 1 (NHE1) to bicarbonate 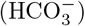 dependent Cl^−^ uptake through a member of the solute carrier family 4 (Novak and Young, 1986; Ogawa et al, 1998; Melvin et al, 2005). For many years, the principal candidate for this role was anion exchanger 2 (Ae2, encoded by the Slc4a2 gene), a member of the solute carrier family 4 (Nauntofte, 1992; Evans et al, 2000; Nguyen et al, 2004; Melvin et al, 2005; Roussa, 2011; Frizzell and Hanrahan, 2012). Direct experimental testing of this hypothesis remained difficult because systemic deletion of the Slc4a2 gene causes severe physiological abnormalities and poor survival in mice, with most dying around the time of weaning (Gawenis et al, 2004, 2010).

Peña-Münzenmayer et al (2015) overcame this limitation by deleting Ae2 specifically in mouse acinar cells, allowing its contribution to salivary secretion to be tested directly. These results challenged the prevailing view. They observed that loss of Ae2 produced little detectable change in stimulated salivary secretion. In contrast, systemic deletion of another member of the same family, anion exchanger 4 (Ae4, encoded by the Slc4a9 gene), reduced secretion by approximately 35% (Peña-Münzenmayer et al, 2015). These experiments also showed that Ae4 knockout acinar cells had a substantially lower resting intracellular Cl^−^ concentration and impaired Cl^−^ uptake during combined CCh and IPR stimulation. Furthermore, the secretory deficit became apparent during sustained secretion rather than as an immediate loss of fluid flow.

In retrospect, these observations already contained important clues to the role of Ae4. The lower resting Cl^−^ concentration suggested that Ae4 influences the chloride available at the onset of secretion, whilst the sustained deficit pointed to a continuing requirement for chloride uptake as secretion proceeds (Peña-Münzenmayer et al, 2015). Our model did not place either observation at the centre of its mechanistic explanation (Vera-Sigüenza et al, 2018). At the time, however, another observation offered a compelling alternative. Subsequent physiological experiments demonstrated that Ae4, unlike Ae2, is an electroneutral, monovalent cation-dependent 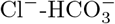 exchanger (Peña-Münzenmayer et al, 2016). Interestingly, this functional characterisation was recently corroborated at the structural level by the identification of the specific molecular determinants governing 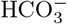 and cation transport within human Ae4 (Catalán et al, 2025). The confirmation of this cation-coupled transport mechanism raised the possibility that the distinctive secretory role of Ae4 arose from its transport of monovalent cations rather than from chloride availability itself. Indeed, by adhering to this point of view, we successfully reproduced the contrasting effects of Ae2 and Ae4 loss, concluding that cation transport was the primary driver of secretion (Vera-Sigüenza et al, 2018).

Experimental work published during and after the development of our original model substantially advanced our understanding of the mechanism by which Ae4 supports salivary secretion. Ae4 was shown to be activated by *β*-adrenergic stimulation through the cAMP-dependent PKA pathway, indicating that its activity is recruited during stimulation rather than functioning solely as a constitutive Cl^−^ uptake pathway (Peña-Münzenmayer et al, 2021). In addition, cAMP-dependent fluid secretion persisted following acinar specific deletion of the gene encoding Tmem16A (also known as anoctamin 1), the principal apical calcium-activated Cl^−^ channel. This provided evidence for an additional stimulated Cl^−^ conductance (Catalán et al, 2015). Together, these findings alter the mechanistic role that can reasonably be assigned to Ae4 during stimulation. If Ae4 activity and apical Cl^−^ conductance are both recruited during secretion, then the ability of Ae4 to sustain intracellular Cl^−^ becomes a plausible determinant of fluid flow rather than a secondary feature of a predominantly cation driven mechanism. This provides a direct reason to reassess the original interpretation (Vera-Sigüenza et al, 2018).

In this light, we re-evaluate the role of Ae4 using an ion conserving dynamical model of the salivary acinar cell. The model incorporates the experimentally observed difference in resting intracellular Cl^−^ between control and Ae4 knockout cells, the *β*-adrenergic activation of Ae4 through the cAMP-dependent PKA pathway, and the coupled ionic, acid base, electrical, and osmotic processes that sustain fluid secretion. We then use the model to test whether the reduction in salivary flow following Ae4 loss can be explained by differences in intracellular Cl^−^ availability at the onset of stimulation and by the continued contribution of Ae4 to Cl^−^ loading during stimulated secretion.

## 2 Model

The model describes ion and water exchange between three compartments: the interstitial space, cytoplasm, and acinar lumen, denoted by the subscripts *e, i*, and *l*, respectively (Fig. 1). Each membrane flux is represented by a submodel describing the associated transporter, channel, or pump. Together, these fluxes determine the ionic composition of the cell and lumen, their volumes, and the rate of fluid secretion (Vera-Sigüenza et al, 2018).

We assume that the concentration of each chemical species is spatially homogeneous within its compartment. Interstitial concentrations are kept constant, corresponding to a cell immersed in an infinite ionic bath. Intracellular and luminal concentrations, however, change as ions and water enter and leave. The model describes the production of primary saliva and its removal from the acinar lumen; subsequent modification by the gland ducts is not included (Sneyd et al, 2021). The complete equations and parameter values are given in Appendices A and C. Here we describe the principal components of the model and the mathematical formulation used to represent each of them.

### 2.1 Model structure and transport laws

Our model retains the compartmental structure and osmotic water transport framework of Vera-Sigüenza et al (2018), but reformulates the ionic and acid base dynamics in terms of conserved amounts and reversible transport processes. Intracellular and luminal ion amounts evolve through membrane transport, whilst separate apical and basolateral membrane potentials are determined from algebraic membrane potential equations. Acid base regulation is represented by tracking total inorganic carbon and alkalinity, from which pH and the individual carbon species are recovered through chemical equilibrium. The model also includes regulated Ae4 activity, basolateral NBCe1-B mediated 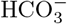 uptake, and an additional stimulated apical Cl^−^ conductance.

For several cotransporters and exchangers, transport is expressed in terms of cycle rates, with the corresponding ionic fluxes determined by their transport stoichiometries. The basolateral NHE1 is represented using the exchanger model of Cha et al (2009), whilst Ae2 and NKCC1 are described by reversible transport laws whose direction and magnitude depend on the prevailing substrate gradients. For NKCC1, the present formulation replaces the reduced expression used previously (Palk et al, 2010). Its driving force depends explicitly on the intracellular and extracellular Na^+^, K^+^, and Cl^−^ concentrations, so net transport vanishes at equilibrium and reverses when the combined gradient changes direction. The complete transport laws are given in Sects. A.5 to A.8.

The model also includes apical K^+^ channels and Na^+^/K^+^ pumps. The apical pump returns K^+^ from the lumen to the cell whilst releasing Na^+^ into the lumen, as proposed in our experimental and modelling study of parotid acinar cells (Almássy et al, 2018). Paracellular 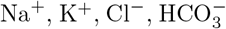, and water transport are represented explicitly. The luminal volume is also dynamic, so water entering the lumen can either leave as secreted fluid or remain temporarily within the acinar lumen. These transport processes are identical in the control and Ae4 knockout cases unless stated otherwise below.

### 2.2 Conserved ionic and acid base state

We denote the amount of species *s* in compartment *r* by *n*_*s,r*_ and its volume by *V*_*r*_. Its concentration is therefore [*s*]_*r*_ = *n*_*s,r*_*/V*_*r*_. A concentration may change through either membrane transport or changes in compartment volume. For instance, the intracellular chloride amount satisfies

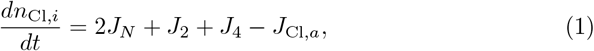

where *J*_*N*_, *J*_2_, and *J*_4_ are the NKCC1, Ae2, and Ae4 cycle rates, respectively. Positive *J*_*N*_, *J*_2_, and *J*_4_ describe Cl^−^ uptake, whilst *J*_Cl,*a*_ denotes total Cl^−^ export through the apical channels. The factor of 2 accounts for the 2 Cl^−^ ions transported during each NKCC1 cycle.

For the acid base system, we assume that the underlying chemical reactions equilibrate rapidly relative to membrane transport. We therefore track the total amounts of inorganic carbon and alkalinity in the cell and lumen, and recover pH and the individual carbon species from the corresponding chemical equilibria. Total inorganic carbon comprises dissolved 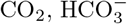, and carbonate. Alkalinity includes 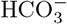, carbonate, the deprotonated buffer, hydroxide, and the negative contribution of free H^+^. The finite buffer can bind or release H^+^, whilst the total number of buffer sites in each compartment remains fixed.

Internal acid base reactions redistribute these species without changing total inorganic carbon or alkalinity. Membrane transport can change either quantity. CO_2_ entry supplies inorganic carbon without supplying alkalinity, whereas 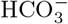 entry supplies both. H^+^ extrusion through NHE1 increases intracellular alkalinity without adding inorganic carbon. Tracking these conserved quantities separately therefore allows the acid base requirements of Cl^−^ transport to be followed without imposing constant pH or an unlimited source of 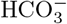.

### 2.3 Acid base requirements of sustained Ae4 transport

Continued Cl^−^ uptake through Ae4 carries an explicit acid base cost. In the transport cycle used here, each positive Ae4 cycle imports one Cl^−^ ion whilst exporting two 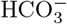 ions (Vera-Sigüenza et al, 2018). Each cycle therefore removes two intracellular alkalinity equivalents. CO_2_ entry can replenish inorganic carbon, but cannot by itself replenish alkalinity, because its hydration produces one 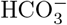 together with one H^+^. Sustained inward Ae4 transport must therefore be accompanied by sufficient alkalinity input through H^+^ extrusion, 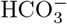 uptake, or another process with the same conserved effect.

Under the transport capacities used in the present model, NHE1 alone does not provide sufficient alkalinity to support the stimulated Ae4 flux whilst maintaining the intracellular state within the imposed physiological range. We therefore include a basolateral 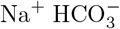 cotransport pathway. Each positive cycle imports one Na^+^ ion together with two 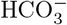 ions, supplying two units of inorganic carbon and two alkalinity equivalents without directly transporting Cl^−^. The intracellular alkalinity equation is

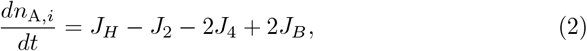

where *J*_*H*_ is the NHE1 cycle rate and *J*_*B*_ is the 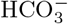 cotransport rate. Positive NHE1 transport increases intracellular alkalinity by extruding H^+^, whilst positive *J*_*B*_ increases it by importing 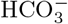. In contrast, Ae2 and Ae4 decrease intracellular alkalinity through 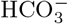 export.

We represent the additional 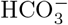 supply using the basolateral electrogenic 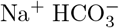 cotransporter NBCe1-B (Slc4a4). NBCe1-B has been identified in mouse submandibular acinar cells, where pNBC1, now termed NBCe1-B, was localised to the basolateral membrane and associated with Na^+^ dependent 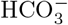 uptake (Luo et al, 2001). The cycle represented here imports one Na^+^ together with two 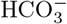 ions, consistent with the electrogenic stoichiometry measured for NBCe1 in salivary acinar cells (Yamaguchi and Ishikawa, 2005). Its reversible flux depends on the Na^+^ and 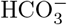 concentrations on both sides of the membrane and on the basolateral membrane potential. Because each inward cycle carries one net negative charge, its charge transport is included in the algebraic equation determining the basolateral membrane potential.

Recruitment of NBCe1-B is prescribed through the Ca^2+^ input, with no additional recruited activity at rest and full recruitment at the reference stimulated Ca^2+^ concentration. This is a modelling assumption introduced to preserve the reconstructed resting state and should not be interpreted as evidence that native NBCe1-B is inactive at rest or regulated by Ca^2+^ in this manner. More generally, the model requires sufficient alkalinity replenishment to offset the 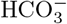 lost through sustained Ae4 transport. NBCe1-B provides a physiologically supported mechanism for this replenishment, but our model does not identify it as the unique pathway.

### 2.4 Ae4 chloride transport and regulation

The Ae4 submodel describes reversible 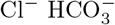 exchange coupled to monovalent cation transport (Peña-Münzenmayer et al, 2016). As in Vera-Sigüenza et al (2018), the retained cycle brings one Cl^−^ ion into the cell whilst exporting two 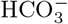 ions and one Na^+^ or potassium ion. We do not remove cation transport from the revised model. Instead, we distinguish the rate of the complete exchange cycle from the allocation of its cation contribution.

The cycle rate is obtained from a carrier model with two membrane facing orientations, a shared Cl^−^ transition, and separate Na^+^ and K^+^ return transitions. Carrier occupancies are assumed to equilibrate rapidly compared with changes in the cellular state. The forward and reverse rates depend on the corresponding substrate gradients. Their construction and the resulting scalar Cl^−^ flux, *J*_4_, are given in the Appendix.

For the principal comparison, the cation contribution is allocated according to the relative Na^+^ and K^+^ concentrations on the side supplying the cations. During Cl^−^ uptake, this is the intracellular side, so the Na^+^ fraction is [Na^+^]_*i*_/[Na^+^]_*i*_ + [K^+^]_*i*_). The remaining fraction is assigned to K^+^. When transport reverses, the extracellular concentrations determine these fractions. This allocation of Ae4 cation transport between Na^+^ and K^+^ is a modelling assumption based on their relative concentrations on the donor side; it is not an experimental measurement of the separate Na^+^ and K^+^ transport rates through Ae4.

To account for the regulatory response demonstrated by Peña-Münzenmayer et al (2021), we introduce an activation variable *x*_4_ that responds to the prescribed *β*-adrenergic input, *β*(*t*). Ae4 activity is multiplied by *m*_4_(*t*), where

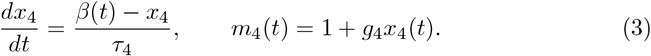

Here *β*(*t*) is zero without stimulation and one during full *β*-adrenergic stimulation. The chosen parameters give an effective activation timescale of 30 seconds and increase the available Ae4 transport activity by up to 25% above its unstimulated value. Eq. (3) represents the net regulatory effect of the cAMP and PKA pathway, rather than modelling cAMP concentration or the individual phosphorylation reactions separately. The magnitude of the gain is consistent with the approximately 25% increase in Ae4 activity reported following forskolin stimulation (Peña-Münzenmayer et al, 2021), although the model’s parameter was not fitted to that particular experiment. Increasing *m*_4_ increases the available transport activity, whilst the evolving ionic state determines the direction and magnitude of Cl^−^ exchange.

### 2.5 Additional stimulated chloride secretion

Vera-Sigüenza et al (2018) allowed apical Cl^−^ secretion only through calcium activated channels. However, the persistence of secretion after Tmem16A knockout under *β*-adrenergic stimulation requires consideration of another route (Catalán et al, 2015). We therefore include an additional apical Cl^−^ conductance alongside Tmem16A. Rather than assigning this conductance to a particular channel protein or prescribing its activation by cell swelling, we represent its response to *β*-adrenergic stimulation by

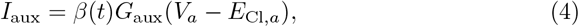

where *G*_aux_ is its conductance, *V*_*a*_ is the intracellular potential relative to the lumen, and *E*_Cl,*a*_ is the apical Cl^−^ equilibrium potential. The current *I*_aux_ enters the algebraic equation determining the apical membrane potential, whilst the corresponding Cl^−^ flux enters the intracellular and luminal Cl^−^ amount equations.

We use the same auxiliary conductance in the control and knockout cases. There is no Ae4 dependent multiplier on either apical Cl^−^ pathway in the principal comparison. The resulting Cl^−^ fluxes can nevertheless differ because the Cl^−^ concentrations and membrane potentials evolve separately in the two cases. Since the conductance of the additional *β*-adrenergic pathway has not been quantified under the conditions represented by the model, we treat *G*_aux_ as an uncertain model parameter and repeat the comparison in its absence and at two nonzero conductance values. The resulting secretion deficit is largely insensitive to these changes, as shown in the Results section.

### 2.6 Control and Ae4 knockout

Removing Ae4 from a cell at the control resting state is not the same experiment as stimulating an established knockout cell. Vera-Sigüenza et al (2018) removed the relevant transporter and allowed the system to approach a new resting state. Here, we use the resting Cl^−^ and pH measurements of Peña-Münzenmayer et al (2015) to define the start of stimulation. The control case begins with [Cl^−^]_*i*_ = 50.10 mM and pH 6.91, whilst the Ae4 knockout case begins with [Cl^−^]_*i*_ = 36.50 mM and pH 6.89.

Changing Cl^−^ alone would disturb both electroneutrality and osmolarity. We therefore retain the Na^+^ concentration and cell volume of the reconstructed control state, together with the fixed intracellular charge and buffer amounts. At the specified Cl^−^ concentration and pH, total inorganic carbon and K^+^ are then calculated to satisfy electroneutrality and equal intracellular and interstitial osmolarity. Alkalinity follows from the acid base equilibria. The lumen and regulatory variables retain their reference initial values. The projection equations and complete initial states are given in Appendix B.

These starting states satisfy charge and osmotic constraints, but are not calculated resting equilibria of the two transport systems. Furthermore, K^+^ and total inorganic carbon differ between them as a consequence of the projection. They are calculated quantities, not additional experimental measurements. The comparison therefore does not isolate Cl^−^ by holding every other intracellular quantity fixed.

Ae4 transport is present in the control case and absent from the knockout. To isolate the effect of the missing chloride supply mediated by Ae4, we prevent differences in NKCC1 and Ae2 activity from compensating for Ae4 loss. At each time point, the NKCC1 and Ae2 cycle rates calculated from the evolving control state are therefore also imposed on the knockout,

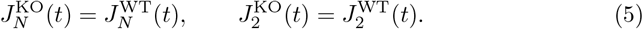

The superscripts WT and KO denote the control and knockout cases. Consequently, the two cases undergo the same NKCC1 and Ae2 transport cycles, and therefore receive the same non Ae4 chloride supply, throughout stimulation, whilst Ae4 mediated chloride uptake remains present only in the control. This is a modelling condition used to isolate the contribution of Ae4 rather than an assumption that NKCC1 and Ae2 are physiologically identical in control and knockout cells. The imposed cycle rates retain their signs and complete transport stoichiometries. They are not held constant in time, but follow the rates calculated from the evolving control state. All remaining transport rates and fluxes are evaluated independently from the state of the corresponding case.

In isolated submandibular acinar cells, Peña-Münzenmayer et al (2015) found no detectable difference in NKCC1 dependent Cl^−^ uptake between Ae4 knockout and control cells when NKCC1 activity was isolated pharmacologically following intra-cellular Cl^−^ depletion. This provides experimental motivation for restricting NKCC1 compensation, although it does not establish equal NKCC1 activity throughout physiological stimulation. No corresponding experimental constraint is available for Ae2 in the Ae4 knockout, so imposing identical Ae2 cycle rates in the two cases is an additional modelling assumption.

### 2.7 Numerical solution

The principal simulations represent combined stimulation with 0.3 *µ*M carbachol (CCh) and 5 *µ*M isoproterenol (IPR) over 600 s, corresponding to the experimental protocol of Peña-Münzenmayer et al (2015). Intracellular Ca^2+^ and the normalised *β*-adrenergic input, *β*(*t*), are prescribed model inputs representing the effects of CCh and IPR, respectively. As in the original Ae4 model, we do not solve the intracellular Ca^2+^ signalling cascade. A detailed model of Ca^2+^ signalling in salivary acinar cells has been developed separately (Pages et al, 2019; Sneyd et al, 2021).

We neglect the fast capacitive dynamics of the membranes and determine the apical and basolateral membrane potentials from algebraic equations at each evaluation of the transport rates. The system is solved numerically using the Radau method with a relative tolerance of 10^−7^ and a maximum time step of 2 s. Further numerical details are given in Appendix D.

Fluid secretion is measured by the luminal outflow, *q*_out_. Because the lumen can change volume, its equation is *dV*_*l*_/*dt* = *q*_*a*_ + *q*_*t*_ −*q*_*out*_, where and are the apical and paracellular water influxes. Cumulative secretion is 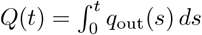, and the relative knockout deficit is 100[1 − *Q*_KO_(*t*)*/Q*_WT_(*t*)].

## 3 Results

Unlike Vera-Sigüenza et al (2018), which examined the new steady states reached following transporter deletion and stimulation, the present simulations follow the experimentally constrained response over the 600 s CCh and IPR stimulation protocol of Peña-Münzenmayer et al (2015). We therefore analyse secretion over this finite experimental interval rather than requiring the system to reach a stationary state. This does not imply that the stationary response is mathematically or physiologically inappropriate; under sustained stimulation the model approaches a slowly varying secretory regime, which can be reproduced and examined directly using the model code available in the GitHub repository (see Appendix G). The finite time analysis is used because it corresponds directly to the duration and initial conditions of the experimental measurement.

### 3.1 Ae4 loss produces a sustained reduction in salivary secretion

Ae4 loss reduced fluid secretion throughout the 600 s stimulation period (Fig. 2). The control produced approximately 0.98 pL of primary fluid, compared with 0.78 pL in the knockout, corresponding to a cumulative reduction of 20.08% calculated from the unrounded volumes. This was smaller than the 35 ± 4.7% reduction reported experimentally by Peña-Münzenmayer et al (2015), but reproduced the direction and substantial magnitude of the secretion deficit. The difference should not be interpreted as a direct failure of the mechanism, since the model represents a single acinar cell and does not reproduce every aspect of the experimental preparation.

**Fig. 2.**
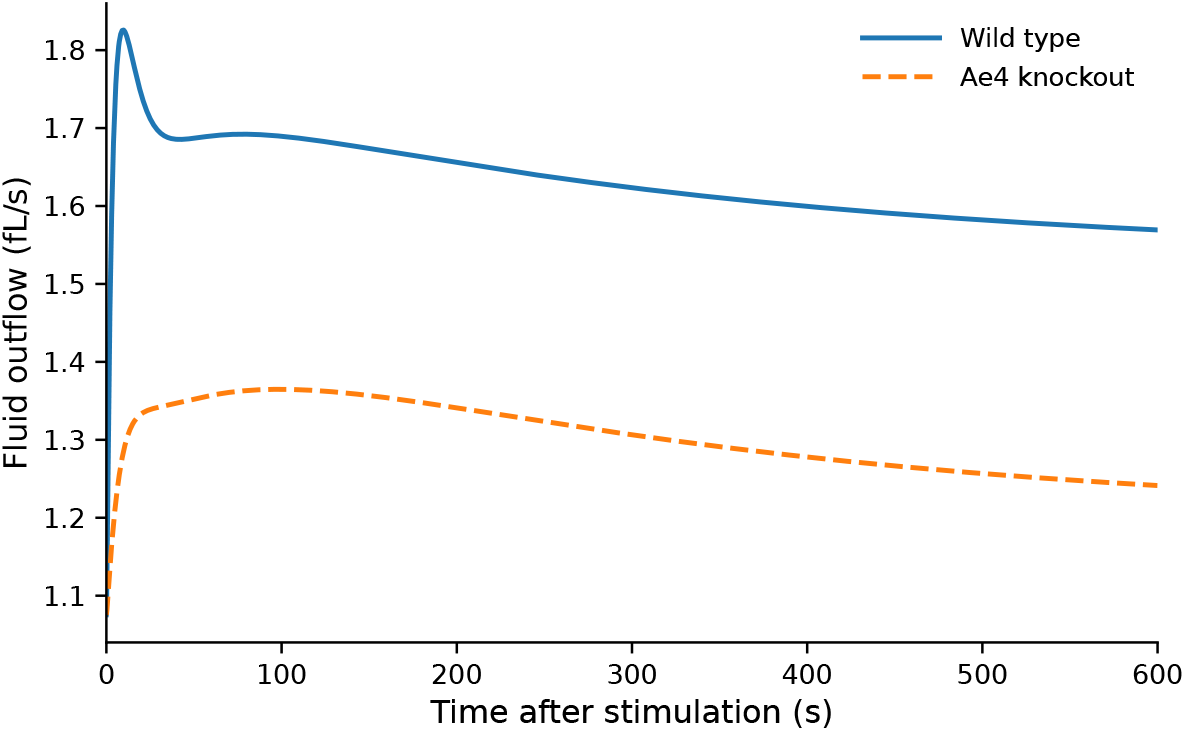
Fluid secretion after Ae4 loss. Luminal outflow from the control and Ae4 knockout cases during combined CCh and IPR stimulation. Ae4 loss reduces the sustained secretion rate throughout the 600 s stimulation period.

The deficit was already present within the first minute and then remained close to 20% for the rest of the protocol. Mean flow between 60 and 600 s was 1.62 fL*/*s in the control and 1.30 fL*/*s in the knockout. This differs somewhat from the experimental response, where the effect of Ae4 loss became more apparent during sustained secretion (Peña-Münzenmayer et al, 2015), but in both cases the reduction persisted throughout prolonged stimulation.

Throughout this period, the control case continued to acquire Cl^−^ through Ae4. Cumulative Ae4 mediated Cl^−^ uptake reached 46.65 fmol over 600 s and remained directed into the cell at the end of stimulation. This source was absent from the knockout, whilst chloride uptake through other transporters remained present. We next examine how this difference affected intracellular Cl^−^ and apical Cl^−^ export.

### 3.2 Lower intracellular chloride is accompanied by reduced apical export

The control and knockout simulations began at intracellular Cl^−^ concentrations of 50.10 and 36.50 mM, respectively, using the experimental means reported by Peña-Münzenmayer et al (2015). Following stimulation, Cl^−^ concentration initially increased in both cases before declining, whilst remaining consistently higher in the control (Fig. 3). At 600 s, intracellular Cl^−^ was 53.71 mM in the control and 37.50 mM in the knockout. Changes in concentration, however, did not directly reflect changes in intracellular Cl^−^ content because cell volume also varied. The control Cl^−^ amount increased from 72.57 to 74.74 fmol, whilst the knockout amount decreased from 52.87 to 50.26 fmol. Both cells contracted during stimulation, reaching final volumes of 1.39 pL in the control and 1.34 pL in the knockout. Consequently, the knockout ended with a slightly higher Cl^−^ concentration than at onset despite containing less intracellular Cl^−^.

**Fig. 3.**
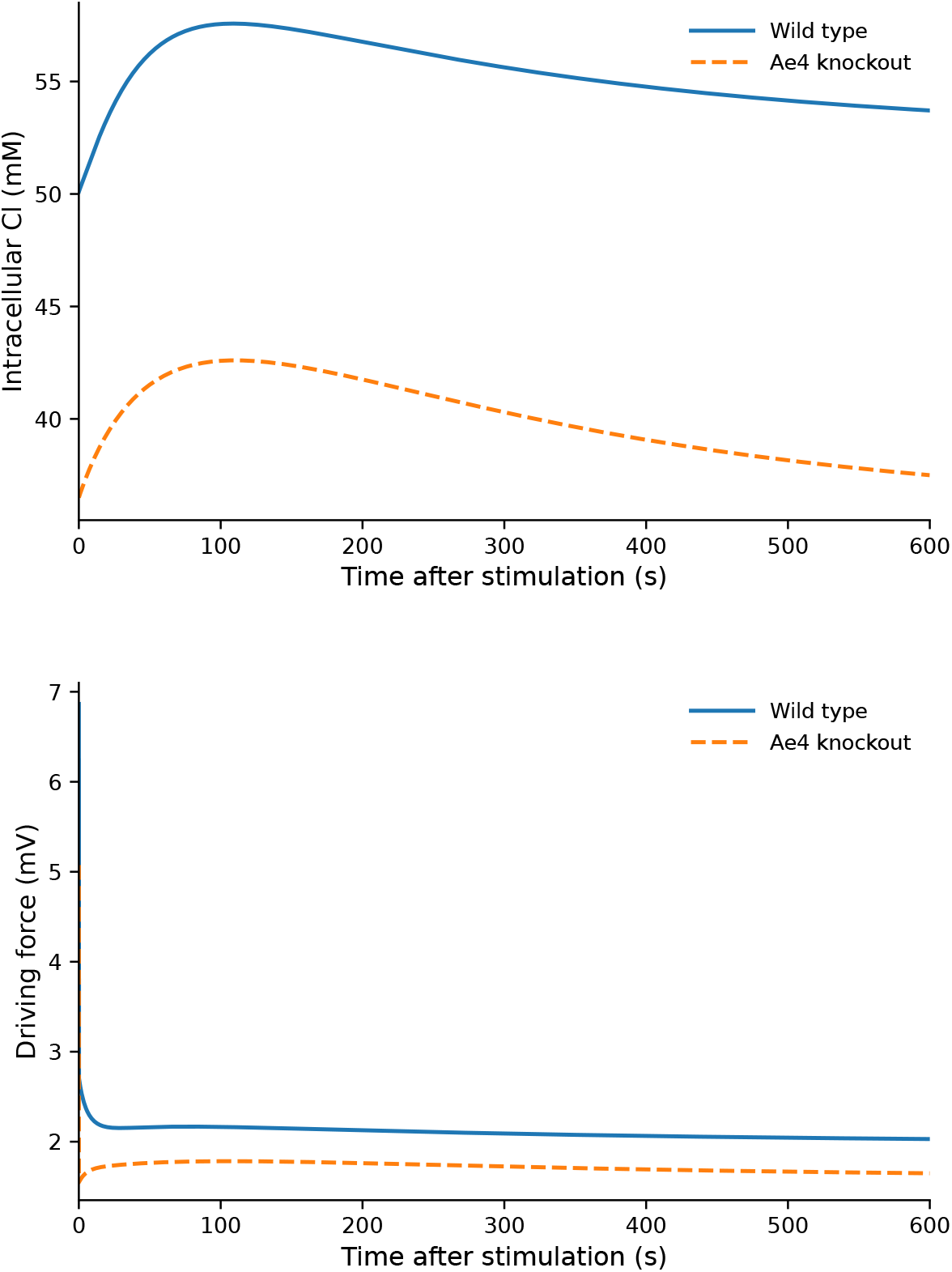
Intracellular chloride and its apical driving force. (A) Intracellular chloride concentrations remain separated throughout combined stimulation. Only the onset concentrations are imposed from the experimental measurements. (B) The outward apical driving force, *E*_Cl,*a*_ − *V*_*a*_, remains positive in both cases but is smaller in the knockout. The chloride channel conductances are identical in the two cases.

This difference in intracellular Cl^−^ altered the apical driving force for secretion. At 600 s, the outward driving force, *E*_Cl,*a*_ − *V*_*a*_, was 2.03 mV in the control and 1.65 mV in the knockout (Fig. 3). Both remained directed outwards, but the weaker driving force in the knockout reduced Cl^−^ export through both apical pathways. At the end of stimulation, total apical Cl^−^ export was 0.37 fmol*/*s in the control and 0.30 fmol*/*s in the knockout, a reduction of 18.75%. This difference arose despite identical apical conductance parameters in the two cases. Over the full simulation, cumulative apical Cl^−^ export was reduced by 18.23% in the knockout, closely matching the 20.08% reduction in cumulative fluid secretion.

The Cl^−^ equation separates the additional apical Cl^−^ export in the control into the initial intracellular Cl^−^ advantage, continued uptake through Ae4, and the intracellular Cl^−^ difference remaining during stimulation. At 600 s, these contributions were 19.70, 46.65, and 24.49 fmol, respectively, giving 41.86 fmol of additional control export. Importantly, the intracellular Cl^−^ difference was larger at the end of stimulation than at its onset. The greater control export therefore cannot be explained by depletion of a larger initial Cl^−^ pool alone; continued Ae4 uptake maintained and further increased the intracellular Cl^−^ advantage throughout stimulation. The explicit acid base formulation also produces a testable prediction. Intracellular pH increased from 6.91 to 7.03 in the control and from 6.89 to 7.27 in the knockout. Experimentally, combined CCh and IPR stimulation also produced intracellular alkalinisation, whilst Peña-Münzenmayer et al (2015) found no significant difference in the initial alkalinisation rate between control and Ae4 knockouts. The larger separation predicted during sustained stimulation therefore points to acid base regulation as an unresolved component of the Ae4 knockout. This is mechanistically important because pH and 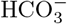 directly influence NHE1, Ae2, Ae4, and NBCe1-B activity, and can therefore feed back on Cl^−^ loading and fluid secretion. The chloride availability mechanism remains supported, but its quantitative magnitude depends partly on the acid base transport processes that regulate intracellular 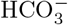, and Cl^−^ loading (Nauntofte, 1992; Melvin et al, 2005; Peña-Münzenmayer et al, 2015). These processes are not yet sufficiently constrained by the available measurements to determine the pH response over longer timescales in control and Ae4 knockout cells.

### 3.3 *β*-adrenergic regulation and the stimulus dependence of secretion

Combined CCh and IPR stimulation recruits Ae4 through the *β*-adrenergic input, representing the cAMP- and PKA-dependent activation demonstrated experimentally by Peña-Münzenmayer et al (2021). Although the prescribed activation increases the available Ae4 transport activity, the realised Cl^−^ flux does not increase monotonically because the ionic driving forces change during secretion. Ae4 mediated Cl^−^ uptake decreased from 0.13 fmol*/*s at stimulation onset to 0.07 fmol*/*s at 600 s, but remained directed into the control cell throughout the protocol. Continued Ae4 uptake therefore contributed to the cumulative Cl^−^ advantage shown in Fig. 4.

**Fig. 4.**
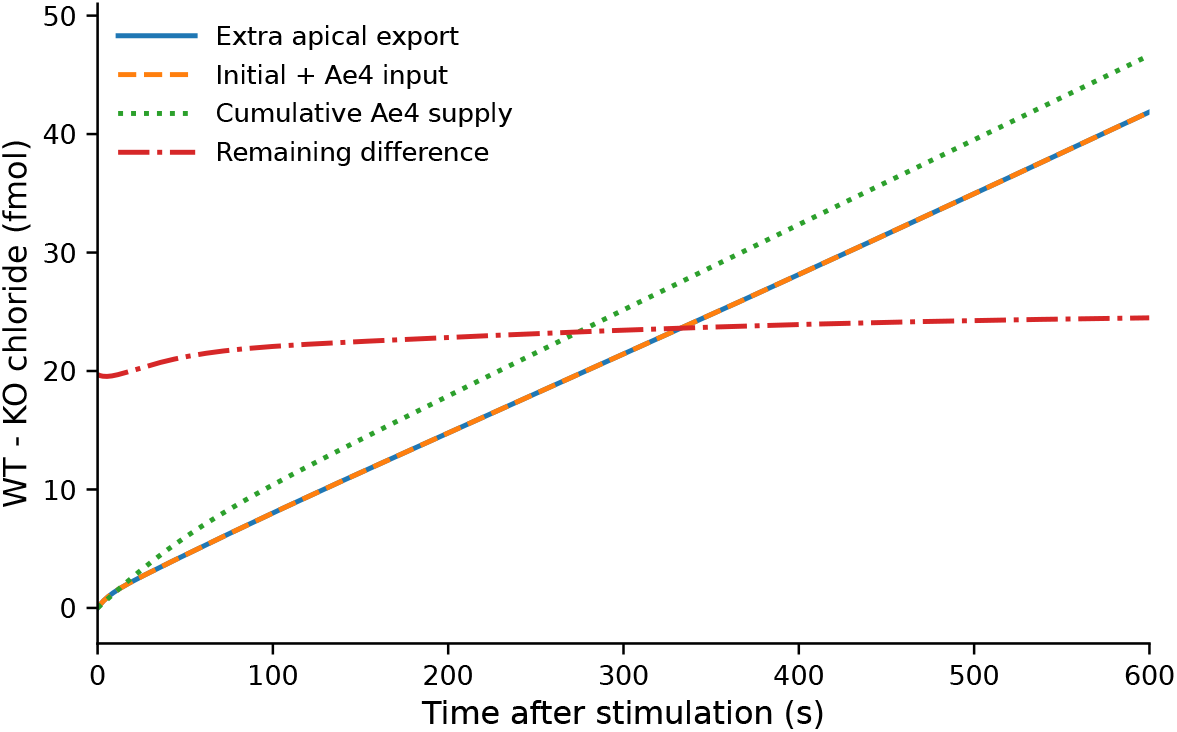
Chloride accounting during combined stimulation. The cumulative excess of control apical Cl^−^ export is accounted for by the initial intracellular Cl^−^ advantage, continued Cl^−^ uptake through Ae4, and the intracellular Cl^−^ difference remaining during stimulation. Ae4 uptake continues throughout the protocol, whilst the intracellular Cl^−^ advantage is maintained rather than depleted.

The additional apical Cl^−^ pathway was also active during combined stimulation. Its cumulative export reached 30.32 fmol in the control and 24.79 fmol in the Ae4 knockout, representing 13.20% of total apical Cl^−^ export in both cases. The conductance was identical in the control and Ae4 knockout; the difference in flux arose from their different Cl^−^ concentrations and membrane potentials.

To examine how the control and Ae4 knockout responses depend on the two stimulus components, we also simulated CCh and IPR separately. CCh alone produced an 18.92% cumulative secretion deficit at 600 s, compared with 20.08% under combined stimulation. Thus, most of the reduction in Ae4 knockout secretion remained in the absence of the *β*-adrenergic input. Under IPR alone, the Ae4 knockout reached the prescribed upper pH limit near 488 s, preventing a valid comparison at 600 s.

The experimental Cl^−^ uptake measurements showed a stronger dependence on stimulus condition. Under CCh alone, the initial uptake rates were 2.18 ± 0.20 in control cells and 2.30 ± 0.10 in Ae4 knockout cells. Under combined CCh and IPR stimulation, the corresponding rates were 2.02 ± 0.10 and 0.90 ± 0.09, whilst IPR alone gave 0.40 ± 0.07 and 0.20 ± 0.03, respectively, in units of 10^−3^ s^−1^ (Peña-Münzenmayer et al, 2015). These experimental values describe initial Cl^−^ recovery, whereas the model comparison describes cumulative fluid secretion over 600 s. They are therefore not directly equivalent observables, but both show that the contribution of Ae4 depends on the stimulation condition.

## 4 Model analysis

The simulations above establish that our model reproduces the principal consequence of Ae4 loss: a decrease in salivary fluid flow. Here we show explicitly how the model equations link intracellular Cl^−^ availability to fluid secretion. In particular, we identify which features of Ae4 distinguish its contribution from the other Cl^−^ uptake pathways within the model.

### 4.1 Intracellular scaling

We use the stimulation duration as the time scale and the initial control cell as the reference for concentration and volume,

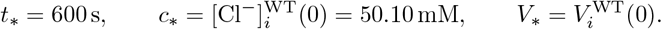

The corresponding amount, transport rate, and water flow scales are

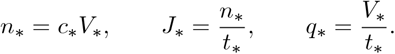

Consequently, transport is measured relative to the initial control Cl^−^ content over one stimulation period. These scales are fixed and shared by the control and Ae4 knockout. For an ionic species *s* in compartment *r*, where *r* = *i* denotes the intra-cellular compartment and *r* = *l* the lumen, let *n*_*s,r*_ denote the corresponding solute amount.

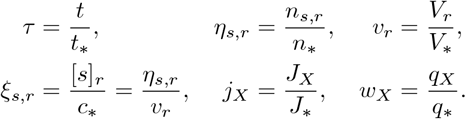

The same amount scale, *n*_∗_, is used to nondimensionalise total inorganic carbon and alkalinity in the cell and lumen. Alkalinity is expressed in equivalents, since the different acid base species contribute different numbers of alkalinity equivalents per mole. Because *J*_∗_ = *n*_∗_ */t*_∗_, the stoichiometric coefficients are preserved under the nondimensionalisation. For example,

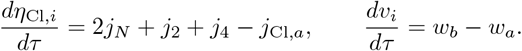

Using the same reference scale for both cases preserves the experimentally imposed difference in resting intracellular Cl^−^ within the dimensionless initial conditions. Membrane potentials are scaled separately by the thermal voltage (Richardson, 2009). Writing *T*_abs_ for absolute temperature,

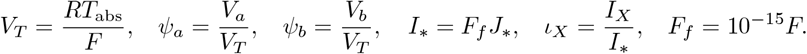

Here *F*_*f*_ converts a monovalent ionic flux measured in fmol s^−1^ into the corresponding electrical current *I*_*X*_. Scaling membrane potential by *V*_*T*_ gives the dimensionless potentials *ψ*_*a*_ and *ψ*_*b*_, which enter the electrochemical driving forces together with logarithmic concentration ratios. Membrane conductances and transporter cycle capacities are nondimensionalised according to their respective units,

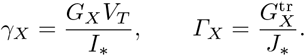

For example,

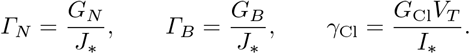

The complete dimensionless system is given in Appendix F.

### 4.2 Chloride availability and apical export

We first separate the effect of the lower initial intracellular Cl^−^ content in the knockout from the effect of continued Ae4 mediated Cl^−^ uptake during stimulation. Under the principal comparison, NKCC1 and Ae2 have the same cycle rates in the control and knockout, whilst Ae4 is present only in the control. Recall that the intracellular Cl^−^ equations are given by

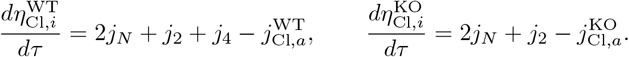

We define the difference in intracellular Cl^−^ amount as 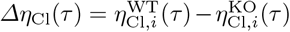. Subtracting the knockout equation from the control equation eliminates the shared NKCC1 and Ae2 contributions,

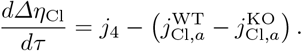

Integrating from the onset of stimulation to time *τ* gives

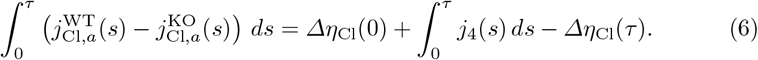

Eq. (6) separates the additional apical Cl^−^ export in the control into the intracellular Cl^−^ difference present at the onset of stimulation, the additional Cl^−^ supplied through Ae4 during stimulation, and the intracellular Cl^−^ difference remaining at time *τ*. This shows that the greater apical Cl^−^ export in the control cannot be attributed solely to its larger initial intracellular Cl^−^ store. If the additional export were supplied only by this initial store, the intracellular difference between the control and Ae4 knockout would decrease during stimulation. Instead, this difference is maintained and increases over the simulated interval. Continued Ae4 mediated Cl^−^ uptake therefore contributes to maintaining the intracellular Cl^−^ available for apical export. Moreover, since 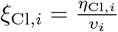, it follows that

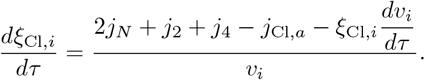

A rise in intracellular Cl^−^ concentration does not therefore imply an increase in intracellular Cl^−^ amount, since cell contraction can increase concentration whilst Cl^−^ is being lost. Eq. (6) is therefore the most appropriate quantity for separating the initial intracellular chloride difference from the additional chloride supplied through Ae4, because it is not confounded by changes in cell volume.

Having separated the contributions of the initial intracellular Cl^−^ difference and continued Ae4 mediated uptake, we next consider how intracellular Cl^−^ availability affects apical export. With the membrane potential scaling defined above, the dimensionless electrochemical driving force for apical Cl^−^ export in case *g* ∈ {WT, KO} is

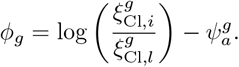

For the prescribed calcium input, let

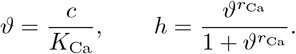

The dimensionless apical Cl^−^ flux is therefore 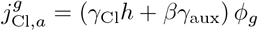. The control and Ae4 knockout have the same apical Cl^−^ conductances and are subjected to the same prescribed Ca^2+^ and *β*-adrenergic inputs. The conductance factor multiplying *ϕ*_*g*_ is therefore identical in the two cases. Whenever this factor is nonzero,

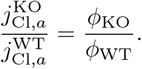

The difference in apical Cl^−^ export therefore arises from differences in the electro-chemical driving force rather than from different channel capacities. The difference between the two driving forces can be written as

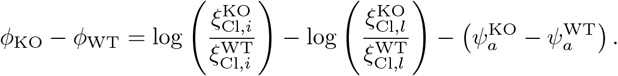

Thus, the difference in intracellular Cl^−^ availability does not determine the difference in apical export on its own. Luminal Cl^−^ and the apical membrane potential also contribute to the electrochemical driving force. The membrane potentials are themselves determined by the transport state. Writing 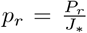, for the pump cycle rates and *ι*_*p*_ for the total paracellular current, the algebraic equations determining the dimensionless membrane potentials are *ι*_K,*a*_ − *j*_Cl,*a*_ + *p*_*a*_ − *ι*_*p*_ = 0, and *ι*_K,*b*_ + *p*_*b*_ + *j*_*B*_ + *ι*_*p*_ = 0. Consequently, the apical membrane potential depends on the coupled transport system rather than on intracellular Cl^−^ alone. In particular, NBCe1-B enters the algebraic equation determining the basolateral membrane potential through *j*_*B*_. It can therefore influence apical Cl^−^ export by altering the membrane potentials as well as through its role in 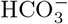 replenishment. For a fixed ionic state and prescribed stimulus, these equations determine a unique pair of membrane potentials. Let *g*_*a*_, *g*_*b*_, and *g*_*p*_ denote the total dimensionless apical, basolateral potassium, and paracellular conductances, respectively. Eliminating *ψ*_*a*_ reduces the two membrane potential equations to a scalar residual *H*(*ψ*_*b*_). Its derivative is

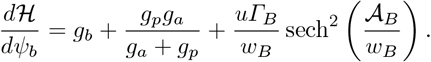

Since the conductances are positive and the final term is nonnegative, 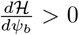. The residual is therefore strictly increasing in *ψ*_b_. Together with its opposite limiting signs as *ψ*_*b*_ tends to positive and negative infinity, this gives a unique solution for the membrane potentials at each fixed ionic state.

Fig. 5 illustrates this result using the control and Ae4 knockout ionic compositions at the end of combined stimulation. The ionic compositions are held fixed whilst *ψ*_*b*_ is varied. The NBCe1-B contribution changes with membrane potential, but the positive channel contribution provides a strictly positive lower bound on *d H /dψ*_*b*_ in both cases. The marked points indicate the membrane potentials obtained from the complete algebraic solution. The effect of intracellular Cl^−^ availability on apical export is therefore mediated by two coupled contributions. Intracellular Cl^−^ enters the electrochemical driving force directly through the concentration ratio, whilst the evolving ionic and transport state determines the membrane potential appearing in the same driving force.

**Fig. 5.**
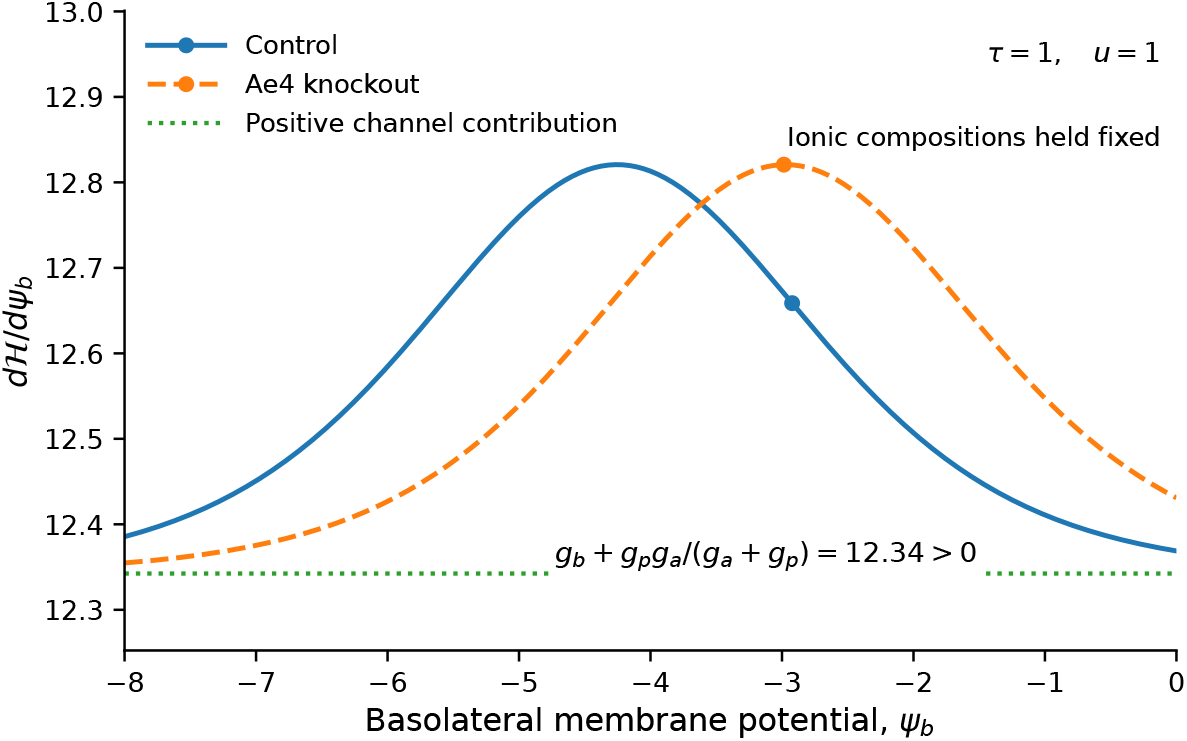
Uniqueness of the algebraic membrane potential solution. The derivative *d H /dψ*_*b*_ is shown as the dimensionless basolateral membrane potential is varied whilst the ionic compositions are fixed at their control and Ae4 knockout values at the end of combined stimulation. The dotted line shows the positive contribution from the channel and paracellular conductances. The derivative remains positive in both cases, consistent with the strict monotonicity of *H*. The marked points indicate the membrane potentials obtained from the complete model state.

**Fig. 6.**
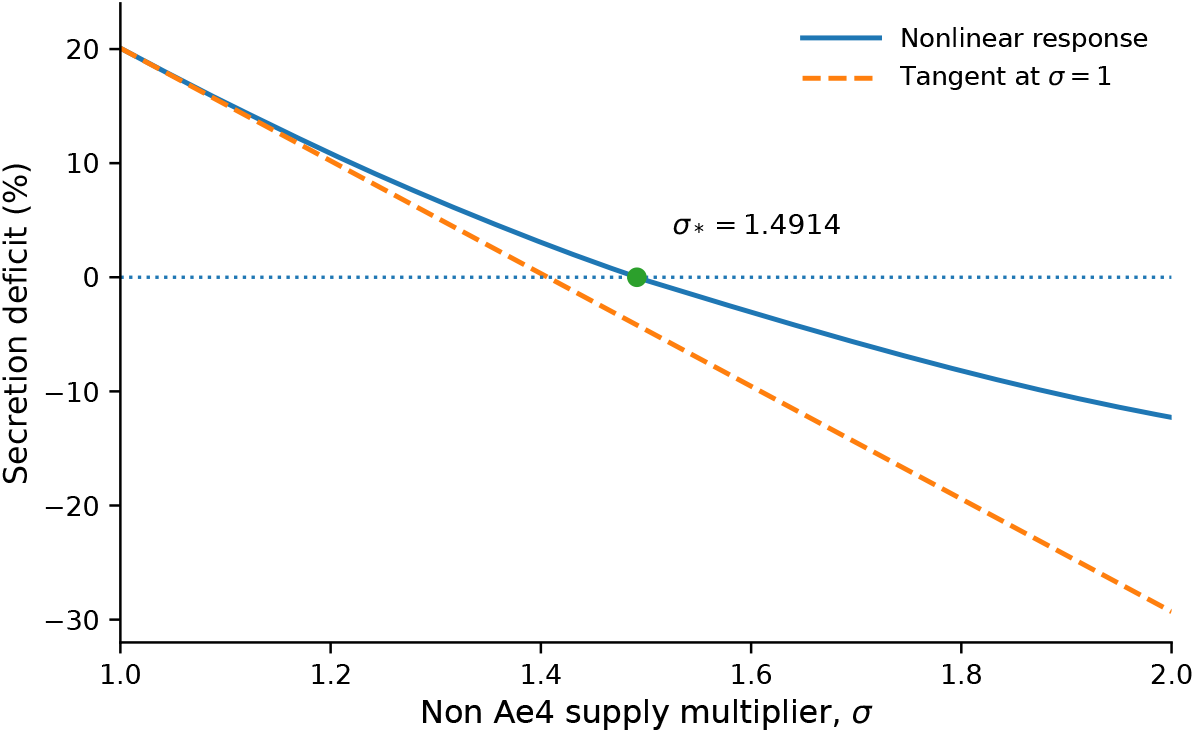
Finite compensation of Ae4 knockout secretion by increased NKCC1 and Ae2 transport. The solid curve shows the nonlinear cumulative secretion deficit as the knockout cycle rates are multiplied by *σ*. The dashed line shows the local approximation about *σ* = 1. Equal cumulative secretion occurs at *σ*_∗_ = 1.49.

### 4.3 Acid base requirements of sustained Ae4 transport

Continued Cl^−^ uptake through Ae4 is coupled to the export of intracellular 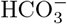. In the transport cycle used here, each positive Ae4 cycle imports one Cl^−^ ion whilst exporting two 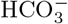 ions and one monovalent cation. Ae4 therefore removes two intracellular alkalinity equivalents per cycle.

In dimensionless form, the intracellular inorganic carbon and alkalinity equations are

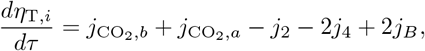

and

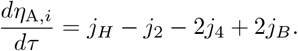

Carbon dioxide entry contributes to total inorganic carbon but not to alkalinity. NHE1 increases intracellular alkalinity through proton extrusion, whilst NBCe1-B supplies both inorganic carbon and alkalinity by importing two 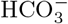 ions per cycle. Rearranging the alkalinity equation and integrating from the onset of stimulation gives

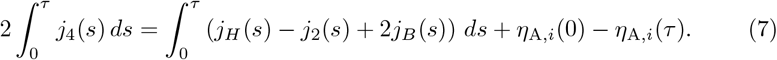

Eq. (7) separates the 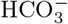 cost of Ae4 transport from the processes that replenish intracellular alkalinity. Over a finite interval, part of this requirement can be supplied by a reduction in the intracellular alkalinity already present at the onset of stimulation. Sustained positive Ae4 transport, however, requires sufficient alkalinity replenishment through proton extrusion, 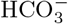 uptake, or another process with the same conserved effect. In the present model, NBCe1-B provides a state dependent 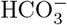 supply, 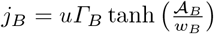, where

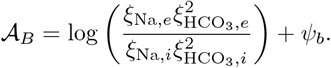

Its contribution therefore depends on both the 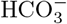 and sodium gradients and the basolateral membrane potential. 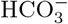 replenishment is not an independent constant source added to the model. It evolves with the cellular state and feeds back on the 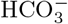 concentration, pH, membrane potential, and transport processes that support continued Cl^−^ loading. The equation in Eq. (7) does not identify NBCe1-B as the unique physiological source of this replenishment. It establishes the more general requirement that sustained Ae4 mediated Cl^−^ uptake must be accompanied by sufficient alkalinity input.

### 4.4 Luminal solute transport and water flow

Apical Cl^−^ export does not determine fluid secretion directly. Water movement depends on the osmotic gradients generated by the combined transport of solutes between the cell, lumen, and extracellular bath. We therefore nondimensionalise osmolarity using the same concentration scale, 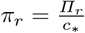, for *r* ∈ {*i, l, e*}. The intracellular and luminal osmolarities are

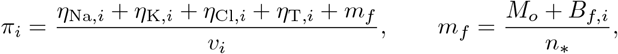

and

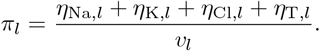

The intracellular osmotic content satisfies

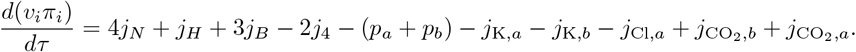

The coefficients follow directly from the transport stoichiometries. One NKCC1 cycle adds four intracellular osmotic particles, one NBCe1-B cycle adds three, whereas one inward Ae4 cycle removes two because the imported Cl^−^ is accompanied by the export of one monovalent cation and two 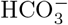 ions. Ae4 therefore does not support secretion by simply increasing intracellular osmolarity. Its contribution depends on how the coupled transport system redistributes solute between the cell and lumen. The dimensionless water fluxes are obtained by defining

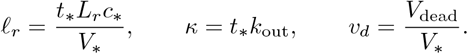

Hence, *w*_*b*_ = ℓ_*b*_(*π*_*i*_ − *π*_*e*_), *w*_*a*_ = ℓ_*a*_(*π*_*l*_ − *π*_*i*_), *w*_*t*_ = ℓ_*t*_(*π*_*l*_ − *π*_*e*_), and *w*_out_ = *κ* max(*v*_*l*_ − *v*_*d*_, 0). The luminal volume therefore satisfies

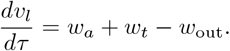

To determine how apical solute transport enters this water equation, we sum the luminal sodium, potassium, chloride, and inorganic carbon equations. This gives

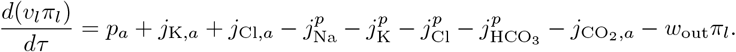

The algebraic equation determining the apical membrane potential is

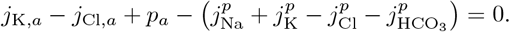

Using this relation to eliminate the cation terms gives the exact equation for luminal osmotic content,

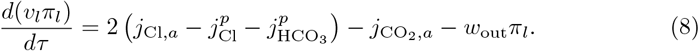

Eq. (8) provides the link between apical Cl^−^ export and the osmotic driving force for water secretion. An increase in apical Cl^−^ export increases luminal osmotic content, but the resulting accumulation also depends on paracellular Cl^−^ and 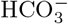 loss, CO_2_ exchange, and removal of solute in the secreted fluid. The factor of two follows from the algebraic equation determining the apical membrane potential and reflects the accompanying net movement of cations required to maintain electroneutrality. Integrating Eq. (8) gives

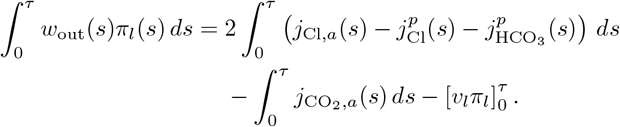

Thus, apical Cl^−^ export and fluid secretion are linked through the accumulation and removal of total luminal solute rather than by a fixed proportionality. A given difference in cumulative Cl^−^ export need not therefore produce the same relative difference in cumulative water secretion.

Finally, cumulative secretion differs from the total water entering the lumen whenever luminal volume changes. Defining 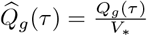, we have

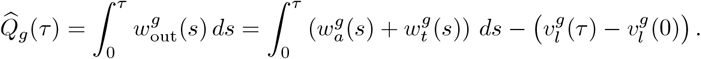

Fluid entering the lumen is therefore either secreted or retained as luminal volume.

### 4.5 Finite compensation by non Ae4 chloride supply

The preceding analysis considers equal NKCC1 and Ae2 cycle rates in the control and Ae4 knockout. We next quantify how much additional non Ae4 transport would be required in the knockout to recover the cumulative fluid secretion of the control. We introduce the dimensionless multiplier *σ* and let 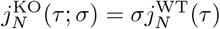, and 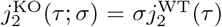. Thus, *σ* = 1 recovers the principal comparison, whilst *σ >* 1 increases the complete NKCC1 and Ae2 transport cycles in the knockout. The control trajectory, the projected initial states, Ae4 expression, and all remaining model parameters are unchanged. The corresponding knockout intracellular Cl^−^ equation is

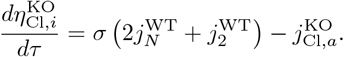

Subtracting this equation from the control equation gives

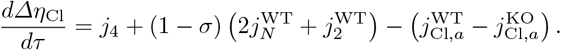

Hence,

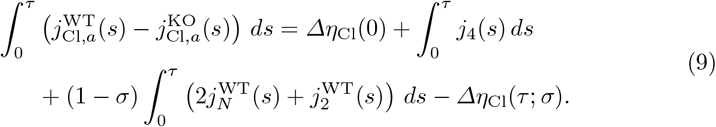

When *σ* = 1, Eq. (9) reduces to Eq. (6). Increasing *σ* adds non Ae4 Cl^−^ supply to the knockout, but the resulting change in apical export also depends on how much of that additional Cl^−^ remains intracellularly. Moreover, because the complete NKCC1 and Ae2 cycles are scaled, the perturbation also changes sodium, potassium, and 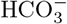 transport. Recovery of fluid secretion therefore cannot be determined by equating the additional Cl^−^ supply with the missing Ae4 flux.

We measure compensation through the cumulative fluid deficit at the end of stimulation,

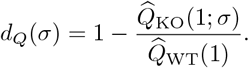

The control trajectory is independent of *σ*, and therefore

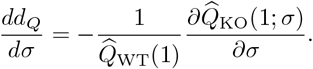

Since

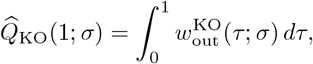

and 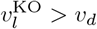 throughout this comparison,

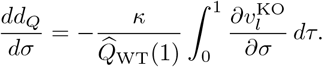

To calculate this derivative, let **z**^KO^(*τ*; *σ*) denote the complete dimensionless knockout state and write its equations as

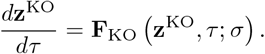

The tangent 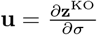 therefore satisfies

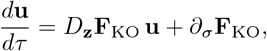

where **u**(0) = 0. The initial condition is zero because the projected knockout onset state does not depend on *σ*. In the state ordering

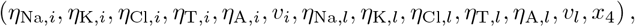

the first five components of *∂*_*σ*_**F**_KO_ are

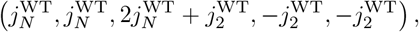

and the remaining eight components are zero. The dependence of pH, the individual carbon species, and the membrane potentials on the evolving knockout state is included when forming *D*_**z**_**F**_KO_; these algebraic quantities are not held fixed in the tangent calculation.

Because 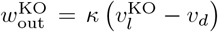 throughout the continuation, the cumulative secretion tangent satisfies

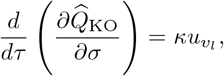

where 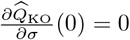, and 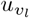 is the luminal volume component of **u**. The corresponding control derivative is identically zero. The local response about the matched supply case is therefore

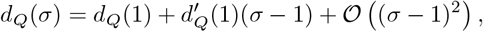

which, for the present model, gives *d*_*Q*_(*σ*) ≃ 0.2 − 0.49(*σ* − 1). Linear extrapolation of this local response predicts equal cumulative secretion at *σ* ≃ 1.41.

We then continued the full nonlinear knockout equations over 1 ≤ *σ* ≤ 2, varying only *σ*. At every value of *σ*, the control retained its original projected onset state, full Ae4 expression, and locally evaluated transport rates, whilst the knockout retained its own projected onset state and zero Ae4 expression.

Natural parameter continuation was used, with the tangent providing the local prediction and nonlinear integration from the unchanged onset state providing the corrected value. The initial parameter step was 0.05 and was halved whenever the discrepancy between the predicted and corrected cumulative secretion deficits exceeded 0.02 percentage points. The accepted continuation contained 41 values of *σ*, separated by 2.50 × 10^−2^. A bracketed scalar solve between the two adjacent continuation values surrounding zero cumulative deficit then gave *σ*_∗_ = 1.49 (rounded to two decimal places). Thus, under this imposed compensation rule, the knockout requires approximately 49.14% greater NKCC1 and Ae2 transport to recover the control cumulative secretion. The local approximation predicts an increase of 40.66%, showing that compensation becomes progressively less effective as the perturbation moves away from the matched supply state.

The value *σ*_∗_ refers only to equality of cumulative model secretion at the end of the stimulation interval. It does not imply equality of the instantaneous secretion rates or of the underlying cellular states. It was not fitted to the experimental Ae4 knockout secretion deficit. The multiplier *σ* is not intended as a physiological model of NKCC1 or Ae2 regulation; it quantifies how strongly these non Ae4 transport cycles must be increased within the present model to compensate for the absence of Ae4. Numerical integration, tangent verification, root calculations, and continuation checks are described in the Appendix.

### 4.6 Charge conservation and chloride export

Intracellular electroneutrality requires *η*_Na,*i*_ + *η*_K,*i*_ − *η*_Cl,*i*_ − *η*_A,*i*_ = *Q*_*f*_ */n*_∗_. The intracellular amounts are therefore constrained and cannot vary independently. This constraint can be combined with the intracellular amount equations to obtain an exact relation for apical Cl^−^ export. Twice the intracellular sodium equation, minus the alkalinity and chloride equations, gives

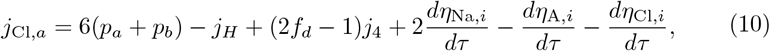

where

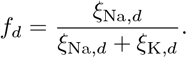

The donor compartment is intracellular during inward Ae4 transport and extracellular when the cycle reverses. NKCC1, Ae2, and NBCe1-B do not appear explicitly in Eq. (10). Their transport rates cancel when the three amount equations are combined. This does not mean that these pathways have no effect on apical Cl^−^ export. They alter intracellular sodium, chloride, alkalinity, pH, and membrane potential, and therefore change the quantities that remain in Eq. (10). The same distinction applies to Ae4. The term (2*f*_*d*_ − 1)*j*_4_ is only its explicit contribution to this particular equation. Removing Ae4 also changes the pump rates, NHE1 transport, and the intracellular accumulation terms. Its total effect on apical Cl^−^ export therefore cannot be inferred from the coefficient multiplying *j*_4_ alone. Eq. (10) also retains the time derivatives of the intracellular amounts. These terms are required because the stimulated trajectories are not assumed to remain at transport equilibrium. Apical Cl^−^ export is therefore determined by both the instantaneous transport rates and the redistribution of intracellular solute during stimulation.

### 4.7 Ae4 carrier parameter identifiability

The Ae4 carrier law contains four microscopic parameters, namely the carrier amount *N*_4_ and the three transition rate scales 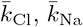, and 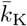. However, these parameters do not enter the whole cell model independently. To make this dependence explicit, choose 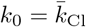, and define

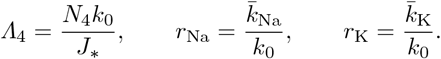

For the Ae4 carrier, define

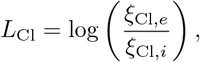

and

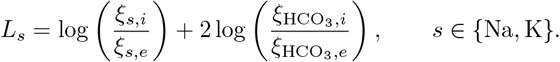

The dimensionless transition rates can then be written as

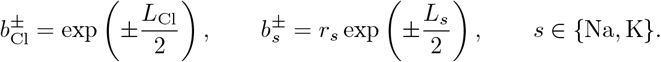

The dimensionless Ae4 cycle rate becomes

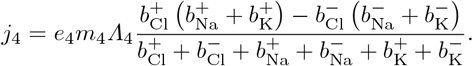

The four carrier parameters therefore enter the cycle rate through only three independent combinations, *Λ*_4_, *r*_Na_, and *r*_K_. This reduction gives an exact scaling symmetry. For any *λ >* 0, the transformation

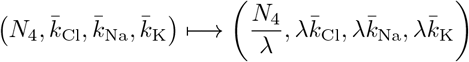

leaves *Λ*_4_, *r*_Na_, and *r*_K_ unchanged. Consequently, *j*_4_ is unchanged at every ionic state. With the same initial conditions and all other model parameters fixed, the complete model trajectory and every derived output, including salivary secretion, are also unchanged. The resulting parameter equivalence is shown in Fig. 7. With *r*_Na_ and *r*_K_ fixed, curves of constant *Λ*_4_ satisfy

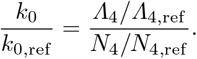

**Fig. 7.**
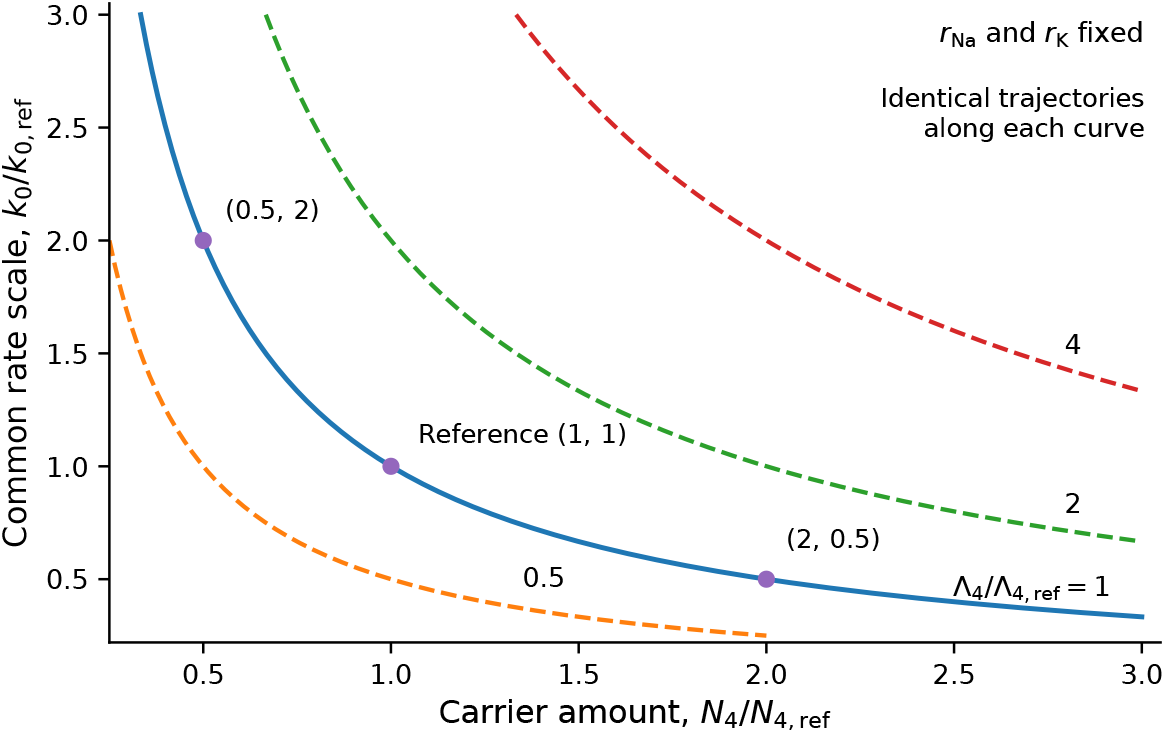
Exact parameter equivalence in the Ae4 carrier model. Curves show constant values of the effective transport combination *Λ*_4_ = *N*_4_*k*_0_*/J*_∗_ with *r*_Na_ and *r*_K_ fixed. The solid curve corresponds to the reference value *Λ*_4_*/Λ*_4,ref_ = 1. The marked parameter pairs lie on this same curve and therefore produce identical Ae4 cycle rates at every ionic state and identical model trajectories when all other parameters and initial conditions are unchanged. Dashed curves show different values of *Λ*_4_.

Thus a reduction in carrier amount can be compensated exactly by a reciprocal increase in the common transition rate scale.

In particular, the points (*N*_4_*/N*_4,ref_, *k*_0_*/k*_0,ref_) = (0.5, 2), (1, 1), and (2, 0.5) lie on the same equivalence curve and therefore generate identical Ae4 cycle rates at every ionic state. Carrier amount and the common transition rate scale are therefore structurally non-identifiable within this formulation. This is not a consequence of limited or noisy secretion data. Even exact observation of the complete model output cannot distinguish an increase in carrier amount from the corresponding decrease in the common transition rate scale. The same symmetry constrains local parameter sensitivities. For a positive parameter *θ*, define

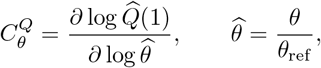

where 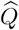 denotes dimensionless cumulative secretion. Differentiating the scaling symmetry with respect to *λ* gives 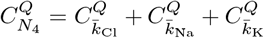. The four local sensitivities are therefore not independent. This limitation persists even if the net Ae4 cycle rate could be reconstructed exactly from the intracellular Cl^−^ equation. Knowledge of *j*_4_(*τ*) would determine the net transport generated by the carrier, but the scaling symmetry would still prevent separate recovery of *N*_4_ and the common transition rate scale. The model can therefore distinguish the effective Ae4 transport combinations that influence secretion, but not the individual microscopic parameters from which those combinations are constructed.

## 5 Discussion

We constructed a model to investigate whether intracellular chloride availability can generate a sustained reduction in fluid secretion following Ae4 loss without requiring any genotype dependent change in apical Cl^−^ conductance.

Experimental evidence implicates Ae4 in this basolateral Cl^−^ loading process. Ae4 knockout reduces resting intracellular Cl^−^, impairs stimulated Cl^−^ uptake, and decreases salivary secretion (Peña-Münzenmayer et al, 2015), whilst subsequent transport measurements established that Ae4 can mediate monovalent cation dependent 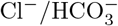 exchange (Peña-Münzenmayer et al, 2016; Catalán et al, 2025). Under the measured onset constraints and prescribed non Ae4 chloride supply used here, the Ae4 knockout secreted 20.08% less fluid than the control over the 600 s stimulation period. The corresponding reduction in cumulative apical Cl^−^ export was 18.23%. Since the apical conductance parameters were identical in the two cases, the reduced secretion arose from differences in the ionic state supplying the shared secretory machinery rather than from an imposed defect in that machinery.

The model shows that the secretion difference cannot be explained solely by the larger intracellular Cl^−^ store present in the control at the onset of stimulation. Continued Cl^−^ uptake through Ae4 also contributes to maintaining the difference during secretion. Experiments have long indicated that sustained secretion requires continued basolateral Cl^−^ uptake to replace Cl^−^ lost through the apical membrane (Nauntofte, 1992; Melvin et al, 2005; Evans et al, 2000). The present simulations make that requirement explicit for Ae4. The control began stimulation with a greater intracellular Cl^−^ amount, consistent with the resting genotype difference measured by Peña-Münzenmayer et al (2015), but it also continued to acquire Cl^−^ through Ae4 throughout. At 600 s, the additional control apical export was accounted for by the initial intracellular Cl^−^ difference, continued Ae4 uptake, and the intracellular difference remaining at the end of stimulation. Importantly, the intracellular Cl^−^ difference was not exhausted during secretion. It was larger at the end of the simulation than at its onset. Ae4 therefore supports secretion by contributing to a sustained difference in intracellular Cl^−^ availability rather than by supplying only a transient reserve. This interpretation is consistent with the broader view that both NKCC1 dependent uptake and parallel 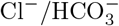 exchange mechanisms contribute to maintenance and recovery of intracellular Cl^−^ during salivary secretion (Nauntofte, 1992; Melvin et al, 2005; Peña-Münzenmayer et al, 2015).

The connection between Cl^−^ availability and secretion is provided by the apical electrochemical driving force. Ca^2+^ dependent activation of apical Cl^−^ conductance is a key component of acinar secretion (Melvin et al, 2005; Sneyd et al, 2021), with Tmem16A providing the principal calcium activated apical Cl^−^ conductance in mouse salivary acinar cells (Catalán et al, 2015). Although the control and knockout share the same apical Cl^−^ conductances in the present model, they do not have the same intracellular Cl^−^ concentration, luminal Cl^−^ concentration, or membrane potential. The resulting outward Cl^−^ driving force is weaker in the knockout, and Cl^−^ export through both Tmem16A and the auxiliary pathway is therefore reduced. Equal channel parameters do not generally produce equal secretory fluxes when the electrochemical states supplying those channels differ. As such, Ae4 acts upstream of the shared apical secretory pathways by altering the Cl^−^ available to them and thereby changing the flux that those pathways can sustain. Nevertheless, this relationship is not a fixed proportionality.

Primary saliva emerges from coordinated ion transport and osmotic water flow rather than from Cl^−^ movement in isolation. Paracellular ion movement and changes in acinar volume also contribute to this process (Nauntofte, 1992; Melvin et al, 2005; Sneyd et al, 2021). Our model analysis reflects this coupling explicitly. Apical Cl^−^ export contributes to luminal osmotic content, but luminal solute accumulation also depends on paracellular Cl^−^ and 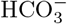 transport, CO_2_ exchange, solute removal in the secreted fluid, and changes in luminal volume. Consequently, a given fractional change in cumulative Cl^−^ export need not generate the same fractional change in cumulative water secretion. The close correspondence between the 18.23% reduction in Cl^−^ export and the 20.08% reduction in fluid secretion in the present simulations is therefore an outcome of the coupled system rather than an imposed conversion between the two quantities.

A second consequence of the model structure is that sustained Ae4 mediated Cl^−^ uptake carries an unavoidable acid base requirement. 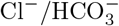 exchange and Na^+^/H^+^ exchange have long been recognised as coupled contributors to Cl^−^ accumulation and intracellular pH regulation in salivary acinar cells (Nauntofte, 1992; Melvin et al, 2005). The Ae4 adds an additional constraint because its transport cycle couples Cl^−^ uptake to the movement of 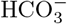 and monovalent cations (Peña-Münzenmayer et al, 2016; Catalán et al, 2025). In the cycle represented here, each positive Ae4 cycle imports one Cl^−^ ion whilst exporting two 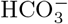 ions and therefore removes two intracellular alkalinity equivalents. Continued Ae4 transport can only be sustained if this loss is offset by sufficient alkalinity input through H^+^ extrusion, 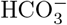 uptake, or another process with the same conserved effect.

In the present model, NBCe1-B provides a source of this alkalinity. Basolateral Na^+^ dependent 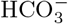 uptake has been demonstrated in mouse submandibular acinar cells, with NBCe1-B localised to the basolateral membrane (Luo et al, 2001). NBCe1-B was omitted from our previous model because it did not affect the predicted secretion under the assumptions used there (Vera-Sigüenza et al, 2018). However, electrogenic Na^+^/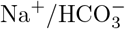 cotransport has also been characterised in salivary acinar cells with a stoichiometry consistent with the transport cycle represented here (Yamaguchi and Ishikawa, 2005). The conclusion of the analysis is nevertheless more general than the particular transporter used in the model. Sustained Ae4 mediated Cl^−^ loading requires coordinated acid base transport, but the present results do not identify NBCe1-B as the unique physiological source of the required alkalinity.

The continuation analysis shows that, under the imposed compensation rule, NKCC1 and Ae2 transport in the knockout must be increased by approximately 49% for cumulative secretion to match the control. NKCC1 is the dominant recognised basolateral Cl^−^ uptake pathway in salivary acinar cells, and both pharmacological and genetic disruption of NKCC1 strongly reduce fluid secretion (Martinez and Cassity, 1983; Evans et al, 2000; Melvin et al, 2005). Nevertheless, secretion persists when NKCC1 activity is absent, supporting the existence of parallel Cl^−^ uptake mechanisms (Evans et al, 2000; Peña-Münzenmayer et al, 2015). In the present model, increasing the complete knockout NKCC1 and Ae2 transport cycles by the common factor *σ* progressively reduced the cumulative secretion deficit. The local response about the principal comparison predicted equal cumulative secretion at *σ* ≃ 1.41, whereas continuation of the full nonlinear system gave *σ*_∗_ = 1.49. Under this prescribed compensation rule, the knockout therefore requires approximately 49% greater NKCC1 and Ae2 transport to recover the control cumulative secretion. This compensation threshold is not a prediction that native NKCC1 or Ae2 regulation produces such an increase. Peña-Münzenmayer et al (2015) found no detectable difference in pharmacologically isolated NKCC1 dependent Cl^−^ uptake between control and Ae4 knockout acinar cells, providing experimental motivation for restricting strong NKCC1 compensation. That experiment does not, however, establish identical NKCC1 flux throughout physiological stimulation, and no equivalent measurement constrains Ae2 under the same conditions. The continuation therefore quantifies how much additional non Ae4 transport would be required within the model rather than identifying an endogenous compensatory response.

The secretion difference is also insensitive to the precise strength assigned to the additional stimulated apical Cl^−^ channel. Acinar specific deletion of Tmem16A does not abolish *β*-adrenergic fluid secretion, demonstrating that an additional secretory pathway can support fluid movement when the principal calcium activated Cl^−^ channel is absent (Catalán et al, 2015). This observation provides the physiological basis for including the auxiliary Cl^−^ conductance in the present model. However, varying that shared conductance over the tested range produced only small changes in the relative control-knockout deficit. The pathway can contribute appreciably to total Cl^−^ export whilst having little effect on the separation between the two cases, because both cases are exposed to the same conductance and their fluxes differ primarily through their electrochemical driving forces. The central Cl^−^ availability mechanism therefore does not depend on tuning an Ae4 specific apical conductance.

In the stimulus specific simulations, most of the predicted secretion deficit remained under CCh stimulation alone, despite Ae4 being activated by *β*-adrenergic cAMP/PKA signalling (Peña-Münzenmayer et al, 2021). The cumulative deficit was 18.92% with CCh alone compared with 20.08% during combined CCh and IPR stimulation, indicating that the effect of Ae4 loss in the model is not primarily dependent on *β*-adrenergic activation. The isolated IPR knockout simulation reached the prescribed pH limit before 600 s, preventing a corresponding sustained comparison under *β*-adrenergic stimulation alone.

The Ae4 carrier formulation also contains an exact parameter ambiguity. The same Ae4 flux can be produced by fewer carriers cycling faster or more carriers cycling more slowly, so whole cell secretion data cannot determine carrier abundance and the common kinetic rate scale separately. Several assumptions define the scope of the model. The resting Cl^−^ and pH values for the control and knockout are imposed from experiment rather than generated from a model of chronic knockout adaptation (Peña-Münzenmayer et al, 2015). Likewise, equal NKCC1 and Ae2 supply in the principal comparison is imposed to isolate the contribution of Ae4. The available NKCC1 data support restricting strong compensation (Peña-Münzenmayer et al, 2015), but neither transporter is sufficiently constrained during sustained secretion to determine how its activity differs between the control and knockout.

Acid base regulation remains the clearest physiological uncertainty. The model explicitly conserves inorganic carbon and alkalinity and therefore exposes the alkalinity demand created by sustained Ae4 transport, but it predicts greater knockout alkalinisation than observed experimentally (Peña-Münzenmayer et al, 2015). The auxiliary apical Cl^−^ pathway is also represented phenomenologically, although the weak sensitivity of the secretion deficit to its conductance shows that the central result does not depend on its precise calibration (Catalán et al, 2015). Finally, the model describes primary secretion from an individual acinar unit, whereas the experimental measurements reflect whole gland output after additional tissue and ductal processes (Melvin et al, 2005; Sneyd et al, 2021). The difference between the predicted 20.08% reduction and the approximately 35% experimental reduction should therefore not be interpreted by subtraction as a missing fraction of the mechanism.

Taken together, the results support a mechanism in which Ae4 maintains intra-cellular Cl^−^ availability both before and during stimulation. This alters the electro-chemical driving force across shared apical Cl^−^ pathways and thereby changes fluid secretion. The same framework also shows that sustained Ae4 transport requires coordinated alkalinity supply and that sufficiently strong non Ae4 Cl^−^ uptake can compensate for its loss.

## A Model equations

The principal comparison considers control and Ae4 knockout cases of the same model. Each cell exchanges ions and water with its own acinar lumen and a common fixed extracellular bath. The only imposed relation between the two cases is the prescribed NKCC1 and Ae2 transport described in Appendix B. Numerical values correspond to the parameter set used throughout this study.

### A.1 Compartments, amounts and units

We denote the interstitial, intracellular, and luminal compartments by *e, i*, and *l*, respectively. For species *s, n*_*s,r*_ is its amount in compartment *r* and *V*_*r*_ is the compartment volume. The concentration is

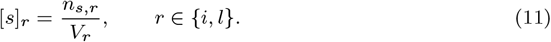

Amounts are expressed in femtomoles (fmol), volumes in picolitres (pL), and concentrations in millimolar (mM). Hence 1 fmol*/*pL = 1 mM. Time is measured in seconds. Calcium inputs and their activation constants are expressed in micromolar (*µ*M).

The state of one cell and its lumen is

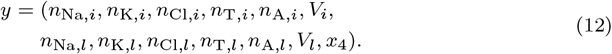

Here *n*_T,*r*_ and *n*_A,*r*_ denote total inorganic carbon and alkalinity amounts, and *x*_4_ is the Ae4 activation variable. Alkalinity amounts have units of fmol equivalents. The state has thirteen differential variables. Membrane potentials and chemical species recovered from equilibrium are algebraic quantities, not additional differential variables.

All transporter cycle rates and ionic fluxes below are whole-cell rates in fmol s^−1^; no additional membrane area or cell volume factor is applied. The individual conventional currents *I*_*s*_ defined below are in amperes and the membrane potentials are in volts. If *I*_*s*_ is associated with ion *s* in a specified direction, its molar flux in the same direction is

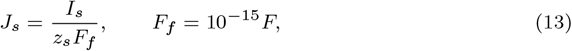

where *z*_*s*_ is the ionic valence and *F* is Faraday’s constant in C mol^−1^. Thus *F*_*f*_ is in C fmol^−1^. For chloride, *z*_*s*_ = −1: an outward chloride flux corresponds to a negative conventional outward current.

### A.2 Carbonate chemistry and finite buffering

Rather than advancing separate kinetic equations for CO_2_, bicarbonate, and H^+^, we recover their concentrations from the total inorganic carbon and alkalinity amounts. This preserves the acid-base chemistry whilst treating the corresponding species as algebraic quantities. Write

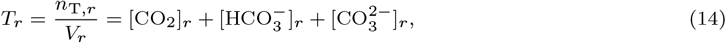

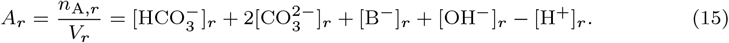

The finite buffer satisfies [BH]_*r*_ + [B^−^]_*r*_ = *B*_*f,r*_*/V*_*r*_, where the site amount *B*_*f,r*_ is fixed.

For the carbonate fractions, H^+^ concentration and dissociation constants are first expressed in molar units,

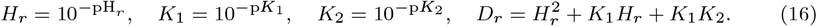

The fractions are

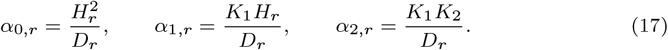

Consequently, the concentrations in mM are

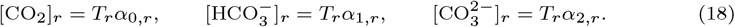

The other equilibrium concentrations, also in mM, are

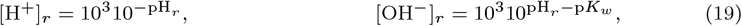

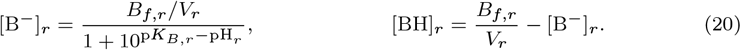

Substitution into Eq. (15) gives one scalar equation for pH_*r*_ at each state evaluation. Neither pH nor bicarbonate is held constant during secretion. The bath uses the same speciation formula with its prescribed pH and total carbon concentration.

Internal hydration and protonation reactions change the distribution of species without changing total inorganic carbon or alkalinity. The relevant external contributions are summarised in Table 1. The buffer site amount does not supply carbon and is counted only once in the intracellular osmolarity.

**Table 1.** Carbon and alkalinity changes per inward molecule or ion. H^+^ extrusion therefore adds one alkalinity equivalent.

| Species entering | Carbon | Alkalinity | Charge |
| --- | --- | --- | --- |
| $CO_2$ | 1 | 0 | 0 |
| $HCO_3^-$ | 1 | 1 | -1 |
| $CO_3^{2-}$ | 1 | 2 | -2 |
| $H^+$ | 0 | -1 | 1 |
| $OH^-$ | 0 | 1 | -1 |

### A.3 Intracellular and luminal equations

Positive *J*_*N*_, *J*_*H*_, *J*_2_, and *J*_*B*_ denote inward NKCC1, NHE1, Ae2, and NBCe1-B cycles. For NHE1, a positive cycle brings Na^+^ in and removes a H^+^; for Ae2, it brings chloride in and removes bicarbonate. The signed Ae4 contributions are denoted 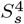. Pump cycle rates *P*_*a*_ and *P*_b_ export Na^+^ across the apical and basolateral membranes. Potassium fluxes *J*_K,*a*_ and *J* _K,*b*_ and total apical chloride flux *J*_Cl,*a*_ are positive out of the cell.

With these conventions, the intracellular equations are

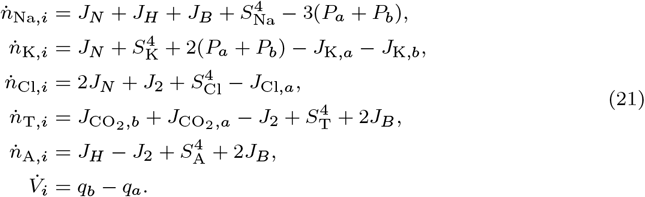

Here 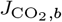 and 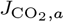 are CO_2_ fluxes from bath and lumen into the cell. The factor of two multiplying *J*_*N*_ in the chloride equation follows from NKCC1 stoichiometry.

Paracellular fluxes 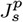 are positive from lumen to bath. The luminal equations are

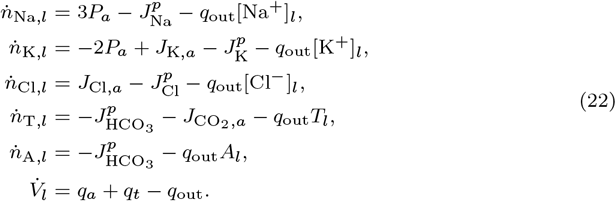

Outflow removes dissolved solutes as well as water. For example, *q*_out_[Cl^−^]_*l*_ has units of fmol s^−1^. The apical pump exports three Na^+^ ions to the lumen and returns two luminal potassium ions to the cell per cycle. Potassium channel and pump localisation are common to both cases, not an imposed knockout defect (Almassy et al, 2018).

The electroneutrality conditions are

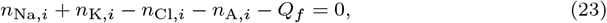

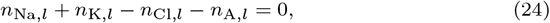

where *Q*_*f*_ is the fixed intracellular anion amount expressed as charge equivalents. This fixed charge is distinct from the buffer contribution already included in alkalinity.

### A.4 Prescribed secretory inputs

Let *c*(*t*) denote intracellular calcium in *µ*M. In the principal combined stimulation protocol,

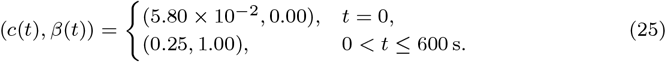

The resting inputs are restored outside the stimulation interval. The protocol represents 0.30 *µ*M carbachol and 5.00 *µ*M isoproterenol, corresponding to the experimental protocol of Peña-Münzenmayer et al (2015). These doses identify the protocol; they do not enter a separately fitted receptor occupancy law. Calcium is prescribed rather than generated by a signalling submodel.

Define the truncation *C*(*v*) = min*{*1, max*{*0, *v}}*. The algebraic recruitment factors are

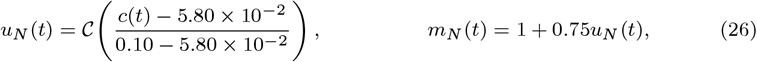

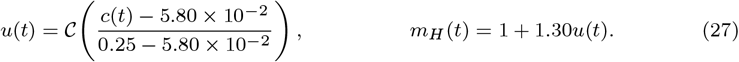

NKCC1 reaches its full multiplier at 0.10 *µ*M, whereas NBCe1-B recruitment and the NHE1 multiplier reach theirs at 0.25 *µ*M. No additional dynamic variable is associated with these factors.

### A.5 NKCC1 and Ae2

The locally evaluated NKCC1 cycle flux is

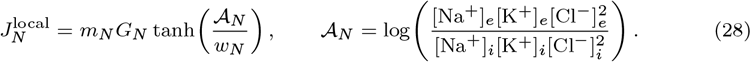

A positive cycle introduces one Na^+^, one potassium, and two chloride ions. The logarithmic concentration ratio determines reversal. The parameter *G*_*N*_ is a cycle capacity rather than a chloride flux capacity. In the principal comparison this local law determines the control cycle rate; the knockout cycle rate is prescribed by Eq. (83).

Ae2 is represented by

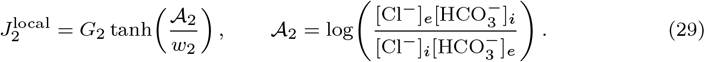

The intracellular source per positive cycle is (0, 0, 1, −1, −1) in the order (Na, K, Cl, T, A). Both equations retain their signs when the concentration gradients reverse; negative rates are not clipped. These bounded reversible laws are effective model choices.

### A.6 Na^+^ H^+^ exchange through NHE1

NHE1 uses the eight state reduction with the second modifier mechanism of Cha et al (2009). All concentrations in the expressions below are in mM. Write

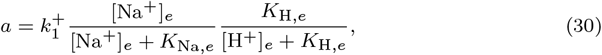

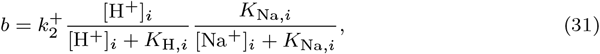

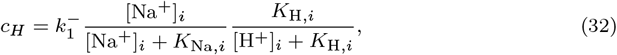

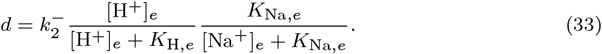

The symbol *c*_*H*_ here is a transition rate, not the prescribed calcium concentration. The modifier and cycle flux are

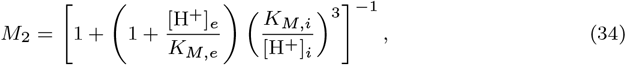

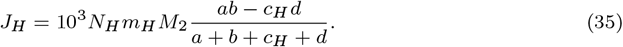

The printed transition rates are in ms^−1^, so the factor 10^3^ converts turnover to s^−1^. *N*_*H*_ is the salivary carrier amount in fmol. The cardiac carrier number in the source study is not transferred to this model. Instead, *N*_*H*_ was selected using the control resting pH reconstruction.

We use the published rounded value 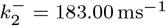. Enforcing the source microscopic reversibility relation on the other rounded constants would give

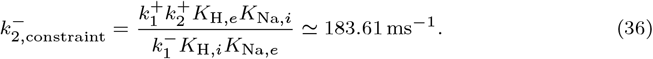

### A.7 NBCe1-B bicarbonate transport and CO_2_ exchange

The basolateral NBCe1-B pathway transports

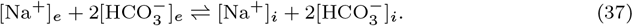

Its inward cycle flux is

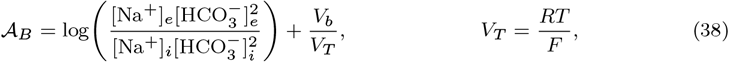

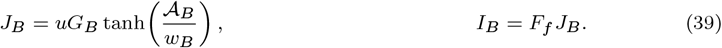

A positive cycle adds one Na^+^, two carbon units, and two alkalinity equivalents to the cell. It imports one net negative charge, giving the positive conventional outward current *I*_*B*_. The dependence on *V*_*b*_ is included when the membrane potentials are calculated. This pathway represents NBCe1-B, which has been localised to the basolateral membrane of mouse submandibular acinar cells and associated with Na^+^ dependent 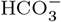 uptake (Luo et al, 2001). The 1 Na^+^ to 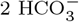 cycle used here is consistent with the electrogenic stoichiometry measured for NBCe1 in salivary acinar cells (Yamaguchi and Ishikawa, 2005).

The capacity *G*_*B*_ was obtained from a control feasibility calculation, not from the knockout secretion endpoint. At the reference chemical state,

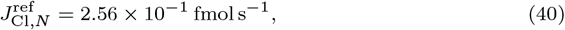

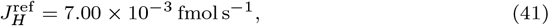

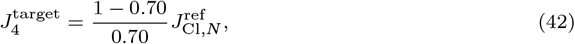

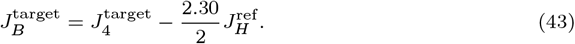

The NBCe1-B flux and electrical equations were then evaluated consistently at the reference chemical state to obtain the capacity in Table 5. The assumed 0.70 fraction sets the scale used to determine *G*_*B*_; it does not constrain the subsequent dynamic flux partition to remain 70*/*30.

CO_2_ crosses both membranes according to

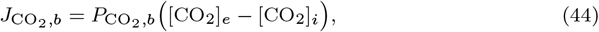

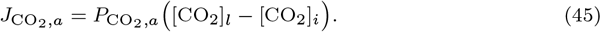

These neutral fluxes enter the total inorganic carbon equations but not the alkalinity equations or the equations determining the membrane potentials. The signs in Eq. (45) are those used in Eq. (21).

### A.8 Ae4 exchange and cation allocation

The scalar Ae4 chloride flux is obtained from a reversible carrier with a common chloride transition and separate Na^+^ and potassium return transitions. For a cation *s* ∈ *{*Na, K}, define

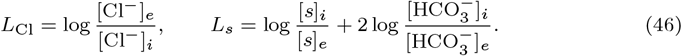

The effective transition rates are

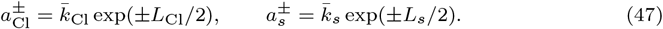

The positive chloride transition proceeds from the outward to inward facing carrier. The positive cation return transitions proceed in the opposite direction. The complete branch affinity is *A*_4,*s*_ = *L*_Cl_ + *L*_*s*_. The rate ratios of this parent carrier therefore recover the corresponding branch affinities.

Let *p*_*o*_ and *p*_*i*_ be the outward and inward facing carrier occupancies. At carrier quasi steady state,

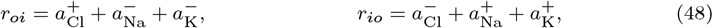

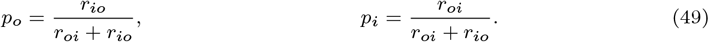

Thus *p*_*o*_ + *p*_*i*_ = 1. The two internal branch turnovers and their total are

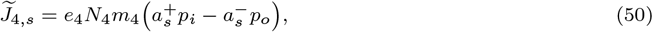

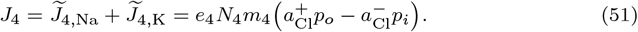

Here *e*_4_ is relative Ae4 expression, *N*_4_ the reference carrier amount, and *m*_4_ the regulatory multiplier. There is no cooperative gate in this parameter set. Cation dependent exchange is supported experimentally, but the carrier amount, attempt rates, and effective cycle representation are model parameters (Peña-Münzenmayer et al, 2016; Catalán et al, 2025).

The principal model uses *J*_4_ to construct its whole-cell sources through a separate donor allocation. Define

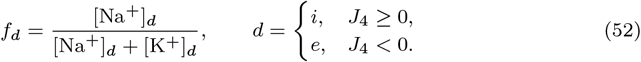

The actual sources used in the amount equations are

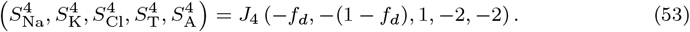

These contributions are electroneutral because 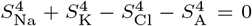. Importantly, the allocated cation sources in Eq. (53) are not identified with the internal branch turnovers in Eq. (51). The source allocation is an effective modelling assumption. The reversibility properties of the parent carrier do not by themselves establish microscopic consistency of this additional allocation.

### A.9 Ae4 activation

The cAMP and PKA response is represented by one effective activation variable,

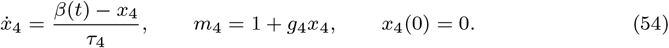

The existence and direction of the regulatory effect follow Peña-Münzenmayer et al (2021). The time constant *τ*_4_ = 30.00 s and gain *g*_4_ = 0.25 are modelling choices rather than measured acute signalling parameters. cAMP, PKA, and phosphorylation states are not modelled individually. In the knockout, *e*_4_ = 0 makes every Ae4 source zero; the regulatory coordinate remains in the paired system.

### A.10 Pump, channels and paracellular currents

The membrane potentials are

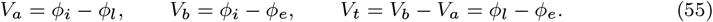

For a species of valence *z*_*s*_, the Nernst potential between compartments *r* and *q* is

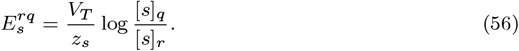

In particular, 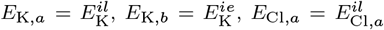, and 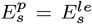. The total Na^+^/K^+^ pump capacity is partitioned between the membranes by *f*_*P*_. Let *P*_*a*,max_ = *f*_*P*_ *P*_max_ and *P*_*b*,max_ = (1 − *f*_*P*_)*P*_max_. The rates are

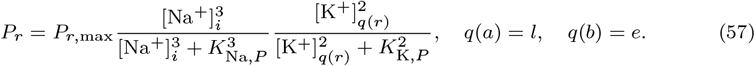

The associated conventional current is *I*_*P,r*_ = *F*_*f*_ *P*_*r*_. ATP is not an additional state or limiting substrate in this pump expression.

The calcium-dependent channel gate is

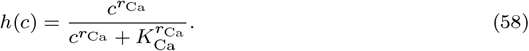

The same gate parameters are used for the potassium and Tmem16A conductances in this parameter set. With potassium partition *f*_*K*_,

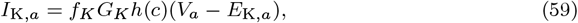

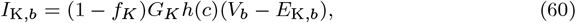

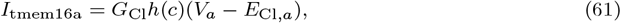

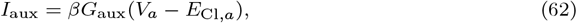

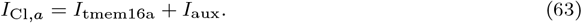

The additional stimulated conductance is motivated by secretion persisting after Tmem16A deletion (Catalán et al, 2015). It is an effective chloride pathway, not an identified channel protein or a resolved swelling mechanism. *G*_aux_ is identical in both cases. The molar chloride flux is *J*_Cl,*a*_ = −*I*_Cl,*a*_*/F*_*f*_.

For *s* ∈ *{*Na, K, Cl, HCO_3_*}*, the paracellular relations are

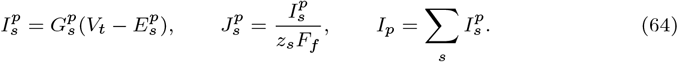

Positive molar flux is from lumen to bath. All four listed paracellular conductances are nonzero in the principal model. They must not be replaced by a Na^+^ and potassium only pathway when reproducing the equations.

### A.11 Membrane potential equations

The fast capacitive dynamics are neglected, so the apical and basolateral membrane potentials are algebraic quantities rather than additional dynamical variables. They are determined at each state evaluation from the individual channel, pump, paracellular, and NBCe1-B currents defined above. Define

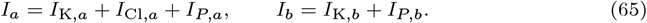

The membrane potentials satisfy

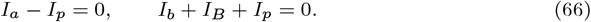

These are algebraic equations used to determine *V*_*a*_ and *V*_*b*_; they do not introduce a separate current state. Since *J*_*B*_, and hence the individual current *I*_*B*_ = *F*_*f*_ *J*_*B*_, depends on *V*_*b*_, NBCe1-B transport and the membrane potentials are evaluated together. For completeness, let

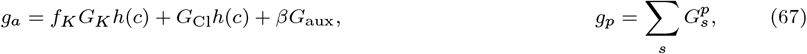

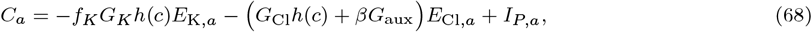

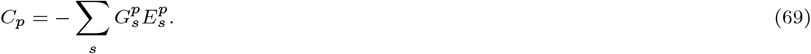

The first membrane potential equation gives

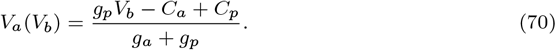

Substitution into the second equation in Eq. (66) leaves a scalar equation for *V*_*b*_. When *u* = 0, *J*_*B*_ = 0 and therefore *I*_*B*_ = 0, so the corresponding linear system for the two membrane potentials is used. These membrane potential equations are solved at every state evaluation, not only at the initial state.

### A.12 Osmotic water flow and luminal outflow

The osmolarities, in mOsm L^−1^, are

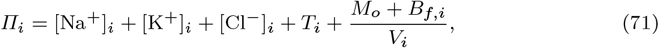

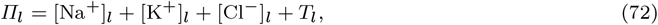

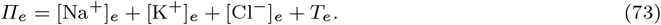

*M*_*o*_ is the amount of intracellular impermeant osmotic particles excluding the finite buffer. Each buffer molecule contributes one osmotic particle regardless of protonation. *Q*_*f*_ is not added again as an osmotic amount. The numerical implementation omits free H^+^ and hydroxide contributions to osmolarity. Its very small luminal buffer amount is included in speciation but not in the luminal osmolarity sum. No additional bath osmolytes are present in this parameter set.

The water fluxes are

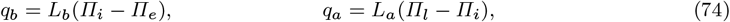

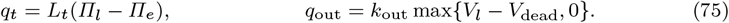

The first three terms are positive from bath to cell, cell to lumen, and bath to lumen, respectively. The final term represents removal of fluid from the finite lumen. It is a prescribed volume-dependent outflow law, not an independently measured ductal pressure relation. In particular,

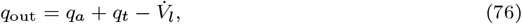

Thus, *q*_out_ = *q*_*a*_ + *q*_*t*_ only when luminal volume is stationary.

## B Initial states and paired comparison

### B.1 Measured onset constraints

The principal comparison imposes [Cl^−^]_*i*_ = 50.10 mM and pH 6.91 in the control case, and [Cl^−^]_*i*_ = 36.50 mM and pH 6.89 in the Ae4 knockout case (Peña-Münzenmayer et al, 2015). These values define the start of stimulation. They are not equilibria calculated from the knockout transport system, and no relaxation is applied before the stimulus. Let *α*_1_ and *α*_2_ be the carbonate fractions at the prescribed pH. Define

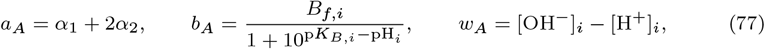

where *b*_*A*_ is an amount and *w*_*A*_ is a concentration. Put *M*_*f*_ = *M*_*o*_ + *B*_*f,i*_. The charge and isotonicity constraints are

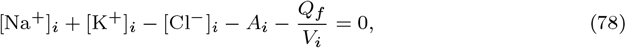

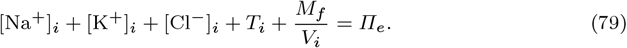

Using *A*_*i*_ = *a*_*A*_*T*_*i*_ + *b*_*A*_*/V*_*i*_ + *w*_*A*_, elimination of potassium yields

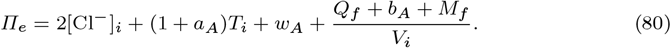

The central projection retains the reference Na^+^ concentration and cell volume,

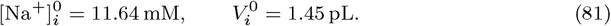

The remaining concentrations are determined without a transport solve,

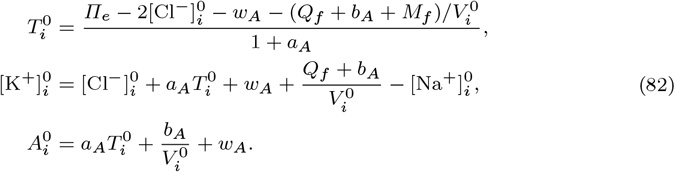

Multiplying concentrations by 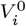 gives the initial amounts. Both cases retain the reference luminal state and 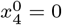. Table 2 gives the resulting complete central states. The larger knockout carbon concentration is a consequence of this projection, not an additional measurement. This is therefore not an isolated chloride perturbation with every other intracellular concentration fixed.

**Table 2.** Central projected onset states. Only the cell chloride concentration and pH are imposed from experiment. Na^+^, volume, and the lumen are inherited; the remaining cell amounts follow from Eq. (82). Displayed numbers are rounded; the accompanying parameter record retains the source precision.

| Coordinate | Control | Ae4 knockout | Units |
| --- | --- | --- | --- |
| $n_{\text{Na},i}$ | 16.85 | 16.85 | fmol |
| $n_{\text{K},i}$ | 168.05 | 166.32 | fmol |
| $n_{\text{Cl},i}$ | 72.57 | 52.87 | fmol |
| $n_{\text{T},i}$ | 24.04 | 45.47 | fmol |
| $n_{\text{A},i}$ | 34.28 | 52.25 | fmol equiv. |
| $V_i$ | 1.45 | 1.45 | pL |
| $n_{\text{Na},l}$ | 13.14 | 13.14 | fmol |
| $n_{\text{K},l}$ | 2.71 | 2.71 | fmol |
| $n_{\text{Cl},l}$ | 15.38 | 15.38 | fmol |
| $n_{\text{T},l}$ | 0.54 | 0.54 | fmol |
| $n_{\text{A},l}$ | 0.47 | 0.47 | fmol equiv. |
| $V_l$ | 0.11 | 0.11 | pL |
| $x_4$ | 0.00 | 0.00 | 1 |

### B.2 Paired supply

Relative Ae4 expression is *e*_4_ = 1 in the control and *e*_4_ = 0 in the knockout. The principal comparison imposes

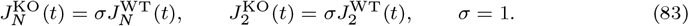

The superscripts denote the control and knockout cases, and the control rates are evaluated from the evolving control state. Each imposed rate retains its sign and complete transport stoichiometry. If Δ*J*_*N*_ and Δ*J*_2_ are the imposed minus local knockout rates, the correction to the knockout’s five intracellular amount derivatives is

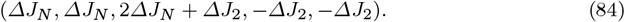

It is electroneutral. No chloride-only source is added and no transporter rate is held constant in time. NHE1, NBCe1-B, pumps, channels, water fluxes, and all algebraic quantities continue to use their respective local states.

The two cases use the same stimulus and *G*_aux_, taking the three values in Table 8. Prescribed rate equality is a modelling condition, not an inference that an isolated NKCC1 uptake assay establishes equality of both physiological fluxes. The finite compensation analysis varies this multiplier over 1.00 ≤ *σ* ≤ 2.00. The single stimulus simulations retain all other parameters and set either *β* = 0 for CCh alone or *c* = 5.80 *×* 10^−2^ *µ*M for IPR alone.

## C Parameter values

Tables 3 to 10 list the numerical parameter values used in the principal model. Where available, parameters were taken from published experimental or modelling studies and are cited accordingly. Remaining values are modelling assumptions or quantities determined from the control case as described in the main text. They should therefore not be interpreted as independent measurements of transporter abundance or activity. Displayed numerical values have at most two decimal places; small quantities and tolerances are written in scientific notation. The numerical simulations use the full precision parameter values in the model implementation.

**Table 3.** Physical and acid base constants. The equilibrium constants are effective values; the model does not apply a separate temperature correction to them. The pH root bracket is a numerical setting, not a physiological range.

| Parameter | Value | Units |
| --- | --- | --- |
| $F$ | 96485.33 | $\text{C mol}^{-1}$ |
| $R$ | 8.31 | $\text{J mol}^{-1}, \text{K}^{-1}$ |
| $T$ | 310.15 | K |
| $\text{p}K_1$ | 6.10 | 1 |
| $\text{p}K_2$ | 10.30 | 1 |
| $\text{p}K_w$ | 14.00 | 1 |
| $\text{p}K_{B,i}$ | 7.00 | 1 |
| $\text{p}K_{B,l}$ | 7.00 | 1 |

**Table 4.** Bath and fixed intracellular quantities. The bath chloride concentration satisfies its charge condition at the stated carbon and pH values. Fixed charge and osmotic amounts are different parameters. The luminal buffer is a near zero model amount, not a measured luminal buffer capacity.

| Parameter | Value | Units |
| --- | --- | --- |
| $[\text{Na}^+]_e$ | 145.00 | mM |
| $[\text{K}^+]_e$ | 5.00 | mM |
| $[\text{Cl}^-]_e$ | 126.16 | mM |
| $T_e$ | 25.00 | mM |
| $\text{pH}_e$ | 7.40 | 1 |
| Bath buffer concentration | 0.00 | mM |
| Additional bath osmolytes | 0.00 | $\text{mOsm L}^{-1}$ |
| $Q_f$ | 78.06 | fmol equiv. |
| $M_o$ | 124.70 | fmol |
| $B_{f,i}$ | 30.00 | fmol |
| $B_{f,l}$ | $1.00 \times 10^{-9}$ | fmol |

**Table 5.**
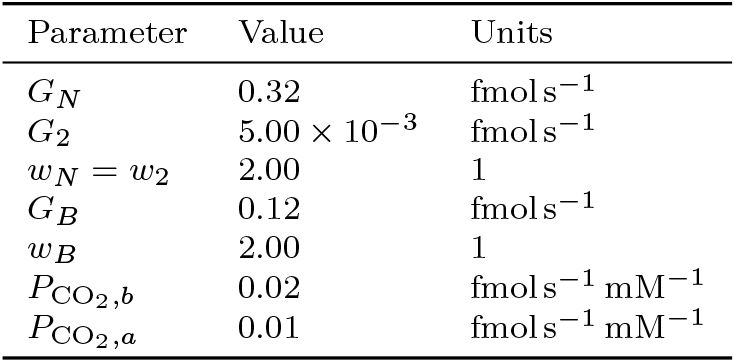
Chloride uptake, bicarbonate supply, and CO_2_ transport parameters. The NBCe1-B capacity is the control feasibility value described in Sect. A.7; it was not selected using the Ae4 knockout secretion deficit.

| Parameter | Value | Units |
| --- | --- | --- |
| $G_N$ | 0.32 | $\text{fmol s}^{-1}$ |
| $G_2$ | $5.00 \times 10^{-3}$ | $\text{fmol s}^{-1}$ |
| $w_N = w_2$ | 2.00 | 1 |
| $G_B$ | 0.12 | $\text{fmol s}^{-1}$ |
| $w_B$ | 2.00 | 1 |
| $P_{\text{CO}_2,b}$ | 0.02 | $\text{fmol s}^{-1} \text{mM}^{-1}$ |
| $P_{\text{CO}_2,a}$ | 0.01 | $\text{fmol s}^{-1} \text{mM}^{-1}$ |

**Table 6.** NHE1 parameters. Transition and binding constants are the published rounded eight state Mod2 values of Cha et al (2009). Only the salivary carrier amount *N*_*H*_ is obtained from the control resting reconstruction. The factor 10^3^ in Eq. (35) is required when the rates are used in the units shown here.

| Parameter | Value | Units |
| --- | --- | --- |
| $N_H$ | $2.34 \times 10^{-5}$ | fmol |
| $k_1^+$ | 10.50 | $\text{ms}^{-1}$ |
| $k_1^-$ | $2.01 \times 10^{-1}$ | $\text{ms}^{-1}$ |
| $k_2^+$ | 15.80 | $\text{ms}^{-1}$ |
| $k_2^-$ | 183.00 | $\text{ms}^{-1}$ |
| $K_{\text{Na},e}$ | 195.00 | mM |
| $K_{\text{Na},i}$ | 16.20 | mM |
| $K_{\text{H},e}$ | $1.62 \times 10^{-3}$ | mM |
| $K_{\text{H},i}$ | $6.05 \times 10^{-4}$ | mM |
| $K_{M,i}$ | $3.07 \times 10^{-5}$ | mM |
| $K_{M,e}$ | $4.80 \times 10^{-7}$ | mM |
| Intracellular modifier exponent | 3 | 1 |
| Extracellular modifier exponent | 1 | 1 |

**Table 7.** Ae4 carrier and regulatory parameters. Carrier amount and attempt rates are model parameters, not measured microscopic constants. The Na^+^ attempt fraction 0.10*/*(0.10 + 1.90) = 0.05 is not the Na^+^ fraction assigned to the physical cation source in Eq. (53).

| Parameter | Value | Units |
| --- | --- | --- |
| $N_4$ | 0.15 | fmol |
| $\bar{k}_{\text{Cl}}$ | 1.00 | $\text{s}^{-1}$ |
| $\bar{k}_{\text{Na}}$ | 0.10 | $\text{s}^{-1}$ |
| $\bar{k}_{\text{K}}$ | 1.90 | $\text{s}^{-1}$ |
| $\tau_4$ | 30.00 | s |
| $g_4$ | 0.25 | 1 |
| $m_4$ at rest | 1.00 | 1 |
| $x_4(0)$ | 0.00 | 1 |
| $e_4$ in control / knockout | 1.00 / 0.00 | 1 |
| Cooperative gate | Absent |  |

**Table 8.**
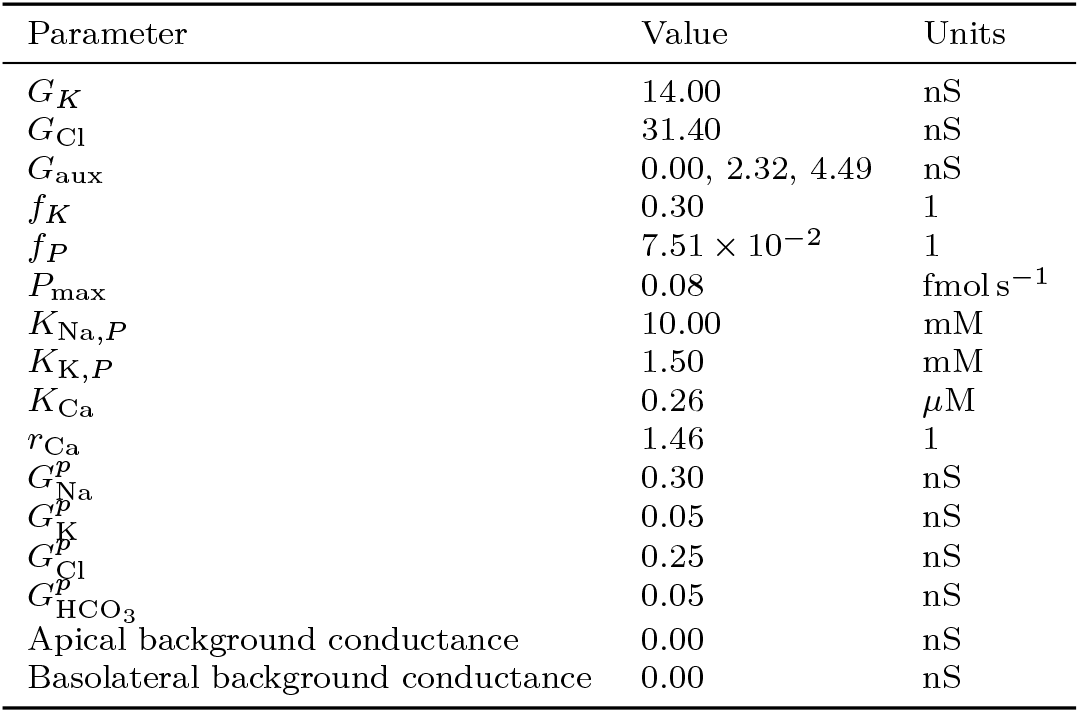
Membrane parameters. Conductances are listed in nS and must be multiplied by 1.00 *×* 10^−9^ when used with voltages in volts to obtain currents in amperes. The same calcium gate is applied to the potassium and Tmem16A conductances. The apical fractions are model choices and should not be interpreted as localisation fractions measured in Almassy et al (2018).

**Table 9.** Water and lumen parameters. The conductances multiply osmolarity differences expressed numerically in mOsm L^−1^. No additional hydrostatic pressure term is included.

| Parameter | Value | Units |
| --- | --- | --- |
| $L_a$ | $4.32 \times 10^{-3}$ | $\text{pL s}^{-1}/(\text{mOsm L}^{-1})$ |
| $L_b$ | $5.15 \times 10^{-2}$ | $\text{pL s}^{-1}/(\text{mOsm L}^{-1})$ |
| $L_t$ | $2.60 \times 10^{-4}$ | $\text{pL s}^{-1}/(\text{mOsm L}^{-1})$ |
| $V_{dead}$ | 0.10 | pL |
| $k_{out}$ | 0.20 | $\text{s}^{-1}$ |

**Table 10.**
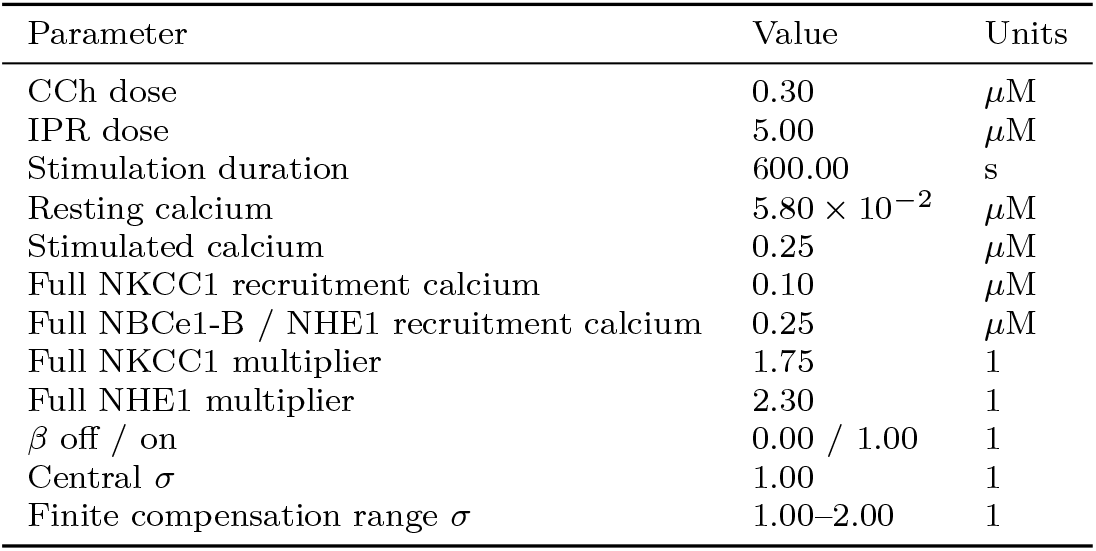
Prescribed stimulation and analysis settings. Agonist doses and duration describe the experimental protocol (Peña-Münzenmayer et al, 2015). Calcium amplitudes, normalised activation endpoints, gains, and supply multipliers are model inputs. A single stimulus simulation changes the indicated input, not the underlying transport parameters.

## D Numerical methods and checks

### D.1 Principal simulations and checks

The principal simulation comprises two copies of the 13-variable state in Eq. (12), with knockout supply prescribed by Eq. (83). It is integrated as a 26-variable system. Carbonate speciation and the two membrane potentials are recovered algebraically whenever its derivatives are evaluated. The numerical settings used for the principal trajectories are given in Table 11.

**Table 11.** Numerical settings for the principal paired system. All tolerances are absolute unless marked relative. The physiological acceptance ranges are separate from the root brackets.

| Setting | Value | Units or interpretation |
| --- | --- | --- |
| Integration method | Radau | implicit method |
| ODE relative tolerance | $1.00 \times 10^{-7}$ | relative |
| Amount absolute tolerance | $1.00 \times 10^{-10}$ | fmol |
| Alkalinity absolute tolerance | $1.00 \times 10^{-10}$ | fmol equiv. |
| Volume absolute tolerance | $1.00 \times 10^{-12}$ | pL |
| Regulatory absolute tolerance | $1.00 \times 10^{-10}$ | fraction |
| Maximum integration step | 2.00 | s |
| Continuous integration start | $1.00 \times 10^{-6}$ | s |
| Reporting interval | 1.00 | s |
| Speciation absolute tolerance | $1.00 \times 10^{-12}$ | pH |
| Speciation relative tolerance | $1.00 \times 10^{-13}$ | relative |
| Voltage absolute tolerance | $1.00 \times 10^{-14}$ | V |
| Voltage relative tolerance | $1.00 \times 10^{-13}$ | relative |
| Projection charge tolerance | $1.00 \times 10^{-9}$ | fmol equiv. |
| Projection isotonicity tolerance | $1.00 \times 10^{-9}$ | mOsm L<sup>-1</sup> |
| Projection pH tolerance | $1.00 \times 10^{-10}$ | pH |
| Independent chloride quadrature | 8 | nodes per interval |
| Quadrature cross check | 4 | nodes per interval |
| Chloride accounting tolerance | $1.00 \times 10^{-5}$ | fmol |
| Quadrature difference tolerance | $1.00 \times 10^{-6}$ | fmol |

At *t* = 0, resting inputs are used. Continuous integration begins at *ϵ* = 1.00 *×* 10^−6^ s from the corresponding projected onset state with stimulated inputs. The onset state is not relaxed or advanced over the omitted interval. The separate zero and right limit samples preserve this convention in the reporting integrals. A BDF fallback is used only if Radau reports solver failure, with unchanged tolerances and inputs. Trajectories that fail the prescribed acceptance checks are not retried with altered parameters.

The pH calculation uses a bracketed scalar solve on [3.00, 11.00]. When NBCe1-B is active, the membrane potential calculation uses the eliminated apical relation and a bracketed solve for *V*_*b*_ on [− 0.50, 0.50] V. These broad numerical brackets are not the acceptance ranges used to classify a physiological trajectory. Root absolute and relative tolerances are stated separately from ODE tolerances.

All accepted solver endpoints and integer-second samples are checked. The first twelve coordinates of each case must be positive, and states, derivatives, and diagnostics must be finite. Additional intracellular acceptance ranges are

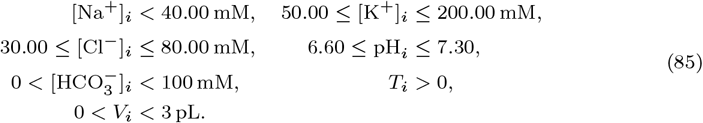

These are the prescribed model checks. They do not amount to a proof of continuous-time validity or long-time stability. A failed trajectory has no accepted endpoint beyond its first detected failed sample.

Residuals of the charge and amount equations are checked in their corresponding units. The residual tolerance is 1.00 *×* 10^−10^ fmol s^−1^ for cell charge rate, lumen charge rate after accounting for outflow, carbon accounting, buffer site accounting, and the electroneutral transporter sources. Water accounting is checked to 1.00 *×* 10^−12^ pL s^−1^, and each algebraic membrane potential equation to 1.00*×*10^−20^ A. The reconstructed alkalinity residual is at most 1.00 *×* 10^−9^ mM in each finite compartment. The auxiliary current components are checked to 1.00 *×* 10^−20^ A; independently assembled cellular source terms and osmotic source terms are checked to 1.00 *×* 10^−10^ fmol s^−1^. The NBCe1-B source is checked against its nonzero physical contribution, − *J*_*B*_ charge equivalents and −2*J*_*B*_ carbon units per second, rather than being incorrectly required to vanish.

Cumulative fluid secretion and the relative deficit are

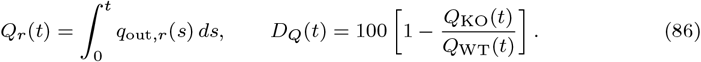

The percentage is defined only when the control integral is positive. For the principal simulations, cumulative quantities are evaluated with the trapezoidal rule on integer-second samples, including the separate zero and *ϵ* samples. Endpoint flow, mean flow over 60 to 600 s, and cumulative flow over 0 to 600 s are distinct observables. No conversion from one model cell to absolute gland volume is applied here.

Independent chloride accounting uses eight-node Gauss–Legendre quadrature on every accepted dense output interval, with four-node quadrature as a cross check. Its absolute chloride accounting tolerance is 1.00 *×* 10^−5^ fmol and its quadrature comparison tolerance is 1.00 *×* 10^−6^ fmol per integrated term. The onset contribution is treated separately, with its flux times duration bound required to satisfy this tolerance. The coarser trapezoidal secretion and export integrals are not used to claim this tighter accounting accuracy.

### D.2 Numerical continuation and verification

The continuation described in Sect. 4.5 is implemented in the dimensional model by varying only the knockout supply multiplier *σ*. The control retains its projected onset state, full Ae4 expression, and locally evaluated transport rates, whilst the knockout retains its own projected onset state and zero Ae4 expression. All remaining parameters and prescribed inputs are unchanged. For each value of *σ*, integration follows the same onset convention as the principal simulations and begins at *t* = 1.00 *×* 10^−6^ s from the corresponding projected onset states. Cumulative secretion is integrated as an additional variable,

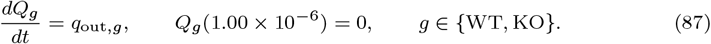

This avoids calculating cumulative secretion from the discrete reporting grid. For the dimensional knockout system,

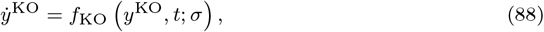

the tangent

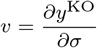

satisfies

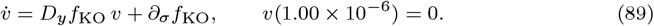

In the state ordering of Eq. (12), the first five components of *∂*_*σ*_*f*_KO_ are

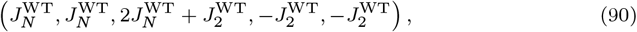

and the remaining eight components are zero. Since *V*_*l*_ *> V*_dead_ throughout the accepted continuation,

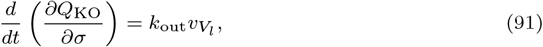

where *v*_*V*_ is the luminal volume component of *v*. The corresponding control derivative is identically zero.

The derivatives in Eq. (89) include the dependence of the recovered chemical species, pH, and membrane potentials on the evolving state. These algebraic quantities are therefore differentiated as part of *D*_*y*_*f*_KO_ rather than held fixed. Their derivatives were evaluated by the complex step method and checked independently against bracketed real root calculations. Across 492 selected cell states, the largest discrepancies were 4.20 *×* 10^−14^ in pH and 1.90 *×* 10^−15^ V in membrane potential.

Natural parameter continuation was performed over

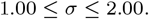

The tangent supplied the local prediction for the cumulative secretion deficit and nonlinear integration from the unchanged onset states supplied the corrected value. The initial parameter step was 0.05. A step was halved whenever the difference between the predicted and corrected deficits exceeded 0.02 percentage points. The accepted continuation contained 41 parameter values separated by 2.50 *×* 10^−2^. A bracketed scalar solve between the adjacent continuation values surrounding zero cumulative deficit gave

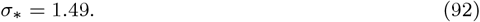

This calculation locates equality of cumulative secretion within the model and does not use the experimental Ae4 knockout secretion deficit as a fitting target.

The trajectories and tangents were integrated with Radau using a relative tolerance of 1.00 *×* 10^−9^ and a maximum time step of 2 s. Absolute tolerances were 1.00 × 10^−11^ for solute amounts and the Ae4 regulatory variable, and 1.00 *×* 10^−13^ for volumes and cumulative secretion. Corresponding absolute tolerances were used for their tangent variables.

The tangent calculation was checked independently using centred perturbations in *σ* at *σ* = 1.00, *σ* = *σ*_∗_, and *σ* = 2.00 over the positive integer-second reporting times. Halving the perturbation approximately quartered the maximum discrepancy, consistent with the expected second-order convergence of the centred difference. At *σ* = 1.00 and *σ* = *σ*_∗_, the discrepancy was below 3.00 × 10^−5^ percentage points per unit *σ* for a perturbation of 2.50 *×* 10^−3^. At *σ* = 2.00, perturbations of 5.00 *×* 10^−3^, 2.50 *×* 10^−3^, and 1.25 *×* 10^−3^ gave a final discrepancy below 7.10 *×* 10^−6^. The wider upper perturbation *σ* = 2.01 reached the prescribed 80.00 mM intracellular chloride limit before 600 s and was therefore not treated as a complete response.

At *σ* = 1.00, *σ* = *σ*_∗_, and *σ* = 2.00, tighter Radau and BDF integrations with relative tolerance 1.00 *×* 10^−10^ and maximum time step 1 s changed the cumulative secretion deficit by less than 1.50 × 10^−9^ percentage points. Independent eight-node flux quadrature gave residuals of the integrated intracellular chloride equation below 4.00 *×* 10^−11^ fmol, with four-node quadrature used as a separate check. Concentration, pH, volume, charge, and algebraic residual checks were applied to accepted integration states and integer-second samples throughout the continuation. These checks establish numerical consistency over the sampled interval 1.00 ≤ *σ* ≤ 2.00; they do not establish validity between the sampled states or beyond this parameter range.

## E Chloride accounting and secretion

Let 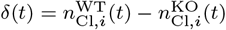 and define the cumulative differences in cycle rates by

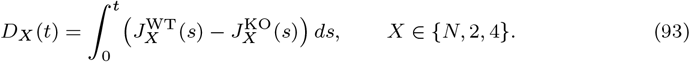

In the principal model, 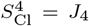. Subtracting the two intracellular chloride equations and integrating gives

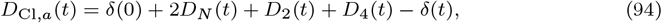

where *D*_Cl,*a*_ is the difference in total apical chloride export through both channels. This identity follows from the amount equations and does not depend on how Ae4 cation transport is allocated.

For the principal comparison, *D*_*N*_ = *D*_2_ = 0 and 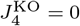, so

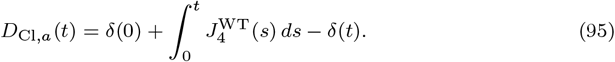

The three terms are the initial control chloride advantage, continued Ae4 input, and the intracellular difference remaining at time *t*. The formula does not assume that either cell exhausts its initial chloride pool.

Nor is apical chloride export identical to chloride removed in saliva. Integrating the luminal chloride equation gives, for either case,

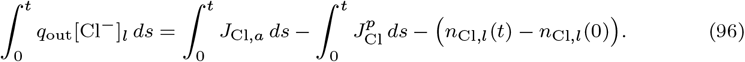

Thus paracellular return and luminal accumulation intervene between apical export and chloride outflow. Water outflow additionally depends on the luminal concentration and osmotic equations. There is no imposed universal conversion factor from a chloride deficit to a fluid deficit.

## F Dimensionless formulation used in the model analysis

The dimensional model is given in Appendix A, and the reference scales are defined in Sect. 4.1. We therefore do not repeat the full transport equations here. Instead, we give the transformations and dimensionless parameter combinations required for the analysis in Sect. 4. All time-dependent inputs are evaluated at *t* = *t*_∗_*τ*.

### F.1 State and algebraic variables

For intracellular and luminal amounts and volumes,

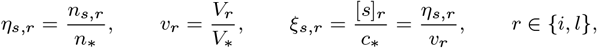

as defined in Sect. 4.1. Fixed extracellular concentrations are scaled in the same way,

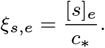

The total inorganic carbon and alkalinity concentrations corresponding to the dimensionless state are

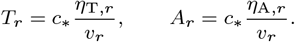

These dimensional concentrations are substituted directly into the carbonate chemistry defined by Eqs. (15) and (17). The resulting equilibrium concentrations are then scaled by *c*_∗_ to obtain, for example,

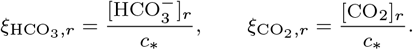

Thus, the nondimensionalisation does not introduce a second acid base model; pH and the individual carbon species are recovered from the same equilibrium equations used in the dimensional system.

The fixed intracellular charge and finite buffer amounts become

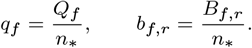

Consequently, Eq. (24) becomes

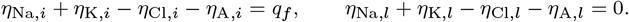

The dimensionless initial conditions are obtained by dividing the amounts and volumes in Table 2 by *n*_∗_ and *V*_∗_, respectively.

### F.2 Transport rates and prescribed inputs

For every transporter cycle or ionic flux,

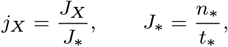

and pump cycle rates are written as

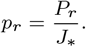

Because *J*_∗_ = *n*_∗_ */t*_∗_, dividing the intracellular and luminal equations in Eqs. (21) and (22) by the corresponding amount or volume scale leaves all transport stoichiometries unchanged. The dimensionless equations used in Sects. 4.2 to 4.4 therefore follow directly from those dimensional equations.

For transport laws with capacities in fmol s^−1^, we define

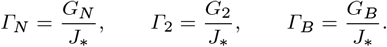

The concentration scale cancels from the logarithmic ratios in Eqs. (28), (29) and (39). Their affinities therefore retain the same form when dimensional concentrations are replaced by *ξ*_*s,r*_. In particular, the NBCe1-B electrical term satisfies

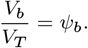

The dimensionless NBCe1-B rate appearing in Sect. 4.3 is consequently obtained directly by dividing Eq. (39) by *J*_∗_.

Similarly, the NHE1 rate is

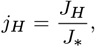

with *J*_*H*_ evaluated from Eq. (35) using the dimensional concentrations [*s*]_*r*_ = *c*_∗_ *ξ*_*s,r*_. No separate NHE1 kinetic law is introduced by the nondimensionalisation.

For CO_2_ transport, division of Eq. (45) by *J*_∗_ gives

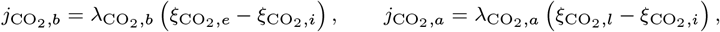

where

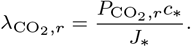

The Ae4 source terms are obtained by dividing Eq. (53) by *J*_∗_. Since all concentrations entering the donor fraction are scaled by the same *c*_∗_,

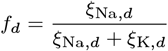

is identical to its dimensional definition. The reduction of the Ae4 carrier law required for the identifiability analysis is given explicitly in Sect. 4.7 and is not repeated here.

The prescribed NKCC1 and Ae2 relation in Eq. (83) likewise becomes

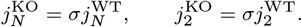

The principal comparison corresponds to *σ* = 1.

Finally, dividing Eq. (54) by the time scale gives

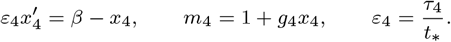

The remaining prescribed calcium-dependent recruitment factors are those of Eq. (27), evaluated at *t* = *t*_∗_*τ*.

### F.3 Membrane potential equations

Membrane potentials are scaled by the thermal voltage and the individual currents by

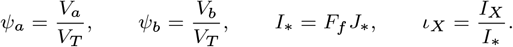

For an ion of valence *z*_*s*_, the Nernst potential in Sect. A.10 therefore becomes

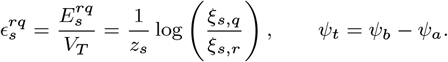

For each channel or paracellular conductance,

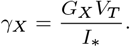

Using the sign convention in Eq. (13), the two algebraic membrane potential equations in Eq. (66) become

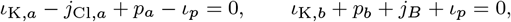

where

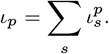

These are the equations used in Sect. 4.2.

To make the scalar membrane potential equation used there explicit, define

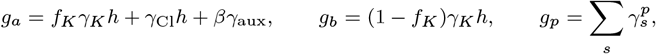

and

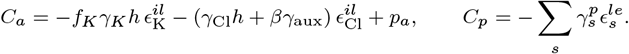

The apical equation then gives

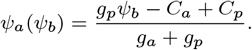

Substitution into the basolateral equation defines

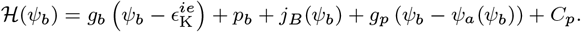

Thus the basolateral membrane potential satisfies

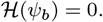

Differentiating this expression gives the derivative used in Sect. 4.2 to establish uniqueness of the membrane potentials for a fixed ionic state and prescribed stimulus.

### F.4 Osmotic variables and secretion

The dimensional osmolarities and water fluxes are given by Eqs. (73) and (75). We define

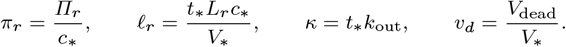

With

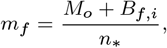

these transformations give the dimensionless osmolarities, water fluxes, and luminal volume equation used in Sect. 4.4. No additional osmotic terms are introduced by the nondimensionalisation.

Finally, cumulative secretion is scaled by the reference cell volume,

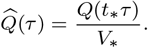

Since *q*_∗_ = *V*_∗_*/t*_∗_,

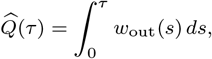

which is the quantity used in the finite compensation analysis of Sect. 4.5.

## G Model implementation and data availability

The implementation of the principal paired model and its numerical records are available in the repository https://github.com/esig626/ae4-salivary-transport-control.

The repository provides the model guide, parameter document, simulation instructions, plotting scripts and an inventory of the accompanying data.

## References

Almássy J, Siguenza E, Skaliczki M, Matesz K, Sneyd J, Yule DI, Nánási PP (2018) New saliva secretion model based on the expression of Na+-K+ pump and K+ channels in the apical membrane of parotid acinar cells. Pflügers Archiv – European Journal of Physiology 470(4):613–621, DOI 10.1007/s00424-018-2109-0

Catalán MA, Kondo Y, Peña-Münzenmayer G, Jaramillo Y, Liu F, Choi S, Crandall E, Borok Z, Flodby P, Shull GE, Melvin JE (2015) A fluid secretion pathway unmasked by acinar-specific Tmem16A gene ablation in the adult mouse salivary gland. Proceedings of the National Academy of Sciences of the United States of America 112(7):2263–2268, DOI 10.1073/pnas.1415739112

Catalán MA, Flores-Aldama L, Fernández F, Bustos D, Apablaza N, Hidalgo A, Mazola Y, Matamala E, Kao LY, Kurtz I, Spichiger C, Sarmiento JM, Brauchi S, González W, Zúñiga L, Peña-Münzenmayer G (2025) Molecular determinants of HCO−3 and cation transport in the human cation-dependent Cl−/HCO−3 exchanger AE4. American Journal of Physiology-Cell Physiology 328(6):C2070–C2084, DOI 10.1152/ajpcell.00346.2024

Cha CY, Oka C, Earm YE, Wakabayashi S, Noma A (2009) A model of Na+/H+ exchanger and its central role in regulation of pH and Na+ in cardiac myocytes. Biophysical Journal 97(10):2674–2683, DOI 10.1016/j.bpj.2009.08.053

Evans RL, Park K, Turner RJ, Watson GE, Nguyen HV, Dennett MR, Hand AR, Flagella M, Shull GE, Melvin JE (2000) Severe impairment of salivation in Na+/K+/2Cl− cotransporter (NKCC1)-deficient mice. Journal of Biological Chemistry 275(35):26,720–26,726, DOI 10.1074/jbc.M003753200

Frizzell R, Hanrahan J (2012) Physiology of epithelial chloride and fluid secretion. Cold Spring Harbor Perspectives in Medicine 2(6):a009.563, DOI 10.1101/cshperspect.a009563

Gawenis LR, Ledoussal C, Judd LM, Prasad V, Alper SL, Stuart-Tilley A, Woo AL, Grisham C, Sanford LP, Doetschman T, Miller ML, Shull GE (2004) Mice with a targeted disruption of the ae2 cl−/hco−3 exchanger are achlorhydric. Journal of Biological Chemistry 279(29):30,531–30,539, DOI 10.1074/jbc.M403779200

Gawenis LR, Bradford EM, Alper SL, Prasad V, Shull GE (2010) Ae2 cl−/hco−3 exchanger is required for normal camp-stimulated anion secretion in murine proximal colon. American Journal of Physiology-Gastrointestinal and Liver Physiology 298(4):G493–G503, DOI 10.1152/ajpgi.00178.2009

Luo X, Choi JY, Ko SB, Pushkin A, Kurtz I, Ahn W, Lee MG, Muallem S (2001) HCO−3 salvage mechanisms in the submandibular gland acinar and duct cells. Journal of Biological Chemistry 276(13):9808–9816, DOI 10.1074/jbc.M008548200

Martinez J, Cassity N (1983) Effect of transport inhibitors on secretion by perfused rat submandibular gland. The American journal of physiology 245(5 Pt 1):G711–6, DOI 10.1016/j.archoralbio.2010.06.012

Melvin JE, Yule D, Shuttleworth T, Begenisich T (2005) Regulation of fluid and electrolyte secretion in salivary gland acinar cells. Annual Review of Physiology 67:445–469, DOI 10.1146/annurev.physiol.67.041703.084745

Nauntofte B (1992) Regulation of electrolyte and fluid secretion in salivary acinar cells. American Journal of Physiology-Gastrointestinal and Liver Physiology 263(6):G823–G837, DOI 0193-1857/92

Nguyen HV, Stuart-Tilley A, Alper SL, Melvin JE (2004) Cl−/HCO−3 exchange is acetazolamide sensitive and activated by a muscarinic receptor-induced [Ca2+]i increase in salivary acinar cells. American Journal of Physiology-Gastrointestinal and Liver Physiology 286(2):G312–G320, DOI 10.1016/j.ceca.2014.01.005

Novak I, Young J (1986) Two independent anion transport systems in rabbit mandibular salivary glands. Pflügers Archiv 407(6):649–656, DOI 10.1007/BF00582647

Ogawa Y, Fernley R, Ito R, Ijuhin N (1998) Immunohistochemistry of carbonic anhydrase isozymes VI and II during development of the rat salivary glands. Histochemistry and Cell Biology 110(1):81–88, DOI 10.1007/s004180050268

Pages N, Vera-Sigüenza E, Rugis J, Kirk V, Yule DI, Sneyd J (2019) A model of Ca2+ dynamics in an accurate reconstruction of parotid acinar cells. Bulletin of Mathematical Biology 81(5):1394–1426, DOI 10.1007/s11538-018-00563-z

Palk L, Sneyd J, Shuttleworth TJ, Yule DI, Crampin EJ (2010) A dynamic model of saliva secretion. Journal of Theoretical Biology 266(4):625–640, DOI 10.1016/j.jtbi.2010.06.027

Peña-Münzenmayer G, Catalán MA, Kondo Y, Jaramillo Y, Liu F, Shull GE, Melvin JE (2015) Ae4 (Slc4a9) anion exchanger drives Cl− uptake-dependent fluid secretion by mouse submandibular gland acinar cells. Journal of Biological Chemistry 290(17):10,677–10,688, DOI 10.1074/jbc.M114.612895

Peña-Münzenmayer G, George AT, Shull GE, Melvin JE, Catalán MA (2016) Ae4 (Slc4a9) is an electroneutral monovalent cation-dependent Cl−/HCO−3 exchanger. Journal of General Physiology 147(5):423–436, DOI 10.1085/jgp.201611571

Peña-Münzenmayer G, Kondo Y, Salinas C, Sarmiento J, Brauchi S, Catalán MA (2021) Activation of the Ae4 (Slc4a9) cation-driven Cl−/HCO−3 exchanger by the cAMP-dependent protein kinase in salivary gland acinar cells. American Journal of Physiology-Gastrointestinal and Liver Physiology 321(6):G628–G638, DOI 10.1152/ajpgi.00145.2021

Richardson G (2009) A multiscale approach to modelling electrochemical processes occurring across the cell membrane with application to transmission of action potentials. Mathematical Medicine and Biology 26(3):201–224, DOI 10.1093/imammb/dqn027

Roussa E (2011) Channels and transporters in salivary glands. Cell and Tissue Research 343(2):263–287, DOI 10.1007/s00441-010-1089-y

Sneyd J, Vera-Sigüenza E, Rugis J, Pages N, Yule DI (2021) Calcium dynamics and water transport in salivary acinar cells. Bulletin of Mathematical Biology 83(4):31, DOI 10.1007/s11538-020-00841-9

Thaysen JH, Thorn NA, Schwartz IL (1954) Excretion of sodium, potassium, chloride and carbon dioxide in human parotid saliva. American Journal of Physiology 178(1):155–159

Vera-Sigüenza E, Catalán MA, Peña-Münzenmayer G, Melvin JE, Sneyd J (2018) A mathematical model supports a key role for Ae4 (Slc4a9) in salivary gland secretion. Bulletin of Mathematical Biology 80(2):255–282, DOI 10.1007/s11538-017-0370-6

Yamaguchi S, Ishikawa T (2005) Electrophysiological characterization of native Na+–HCO−3 cotransporter current in bovine parotid acinar cells. The Journal of Physiology 568(1):181–197, DOI 10.1113/jphysiol.2005.088633

Young J, Schögel E (1966) Micropuncture investigation of sodium and potassium excretion in rat submaxillary saliva. Pflüger’s Archiv für die gesamte Physiologie des Menschen und der Tiere 291(1):85–98, DOI 10.1007/BF00362654

Yusuke I, Hiroshi N C, Yoshizaki M, Gen P (1973) Evidence for the osmotic flow across dog submaxillary gland epithelia as a cause of salivary secretion. The Japanese Journal of Physiology 23(6):635–644, DOI 10.2170/jjphysiol.23.635

